# Live imaging of hair bundle polarity acquisition in the mouse utricle demonstrates a critical timeline for transcription factor Emx2

**DOI:** 10.1101/2020.05.28.121244

**Authors:** Yosuke Tona, Doris K. Wu

**Affiliations:** National Institute on Deafness and Other Communication Disorders, National Institutes of Health, Bethesda, MD 20892, USA

## Abstract

The asymmetric hair bundle on top of hair cells (HCs), comprises a kinocilium and stereocilia staircase, dictates HC directional sensitivity. The mother centriole (MC) forms the base of the kinocilium, where stereocilia are subsequently built next to it. Previously we showed that transcription factor Emx2 reverses hair bundle orientation and its expression in the mouse vestibular utricle is restricted, resulting in two regions of opposite bundle orientation (Jiang et al, 2017). Here, we investigated establishment of opposite bundle orientation in embryonic utricles by live-imaging GFP-labeled centrioles in HCs. The daughter centriole invariably migrated ahead of the MC from the center to their respective peripheral locations in HCs. Comparing HCs between utricular regions, centriole trajectories were similar but they migrated towards opposite directions, suggesting that Emx2 pre-patterned HCs prior to centriole migration. Ectopic Emx2, however, reversed centriole trajectory within hours during a critical time-window when centriole trajectory was responsive to Emx2.

## INTRODUCTION

The mammalian inner ear comprises six major sensory organs including the cochlea, two maculae and three cristae. The cochlea detects sound, whereas the maculae and cristae detect linear accelerations and angular velocity of head movements, respectively. Each sensory organ consists of sensory hair cells (HCs), which are the mechano-transducers of sound and head movements, and each HC is surrounded by supporting cells. Erected on the apical surface of HCs is the stereociliary bundle/hair bundle, which is comprised of a stereociliary staircase that is tethered to the tallest rod of the bundle, the kinocilium. When the hair bundle is deflected towards the kinocilium, the mechano-transducer channels on the tips of the stereocilia open, which allow entry of positive ions and activation of the HC (Shotwell et al., 1981). Thus, orientation of the hair bundle provides the directional sensitivity of its HC.

Each sensory organ of the inner ear exhibits a defined hair bundle orientation pattern among HCs. Unlike other sensory organs in which hair bundles are unidirectional, the macula of the utricle and saccule exhibit an opposite bundle orientation pattern. Each macula can be divided by a line of polarity reversal (LPR) into two regions, across which the hair bundles are arranged in opposite orientations (Fig. 1A; (Flock, 1964)). Although proper alignment of the hair bundles in sensory organs requires the Wnt signaling pathway and the core planar cell polarity pathway (Dabdoub et al., 2003, Goodrich and Strutt, 2011, Jones et al., 2014, Lee et al., 2012, Montcouquiol et al., 2003, Montcouquiol et al., 2006), the LPR in the maculae is generated by the transcription factor Emx2. The restricted expression of Emx2 to one side of the LPR causes hair bundles within to reverse their orientation by 180 degrees (Fig. 1A, green color; (Jiang et al., 2017, Holley et al., 2010)). In *Emx2* knockout or Gain-of-function utricles, the LPR is absent and all hair bundles are unidirectional (Fig. 1B).

**Figure 1.**
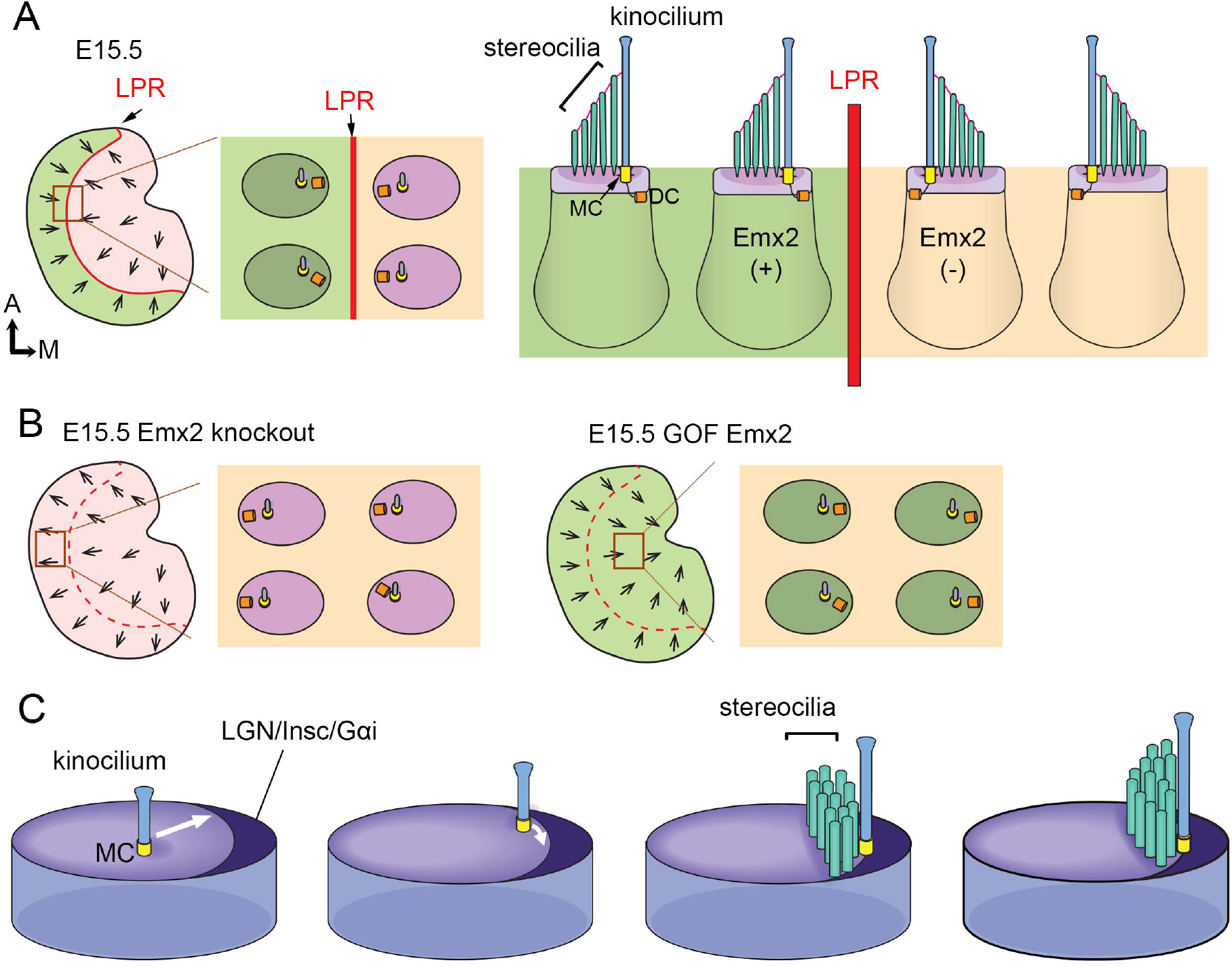
Hair bundle orientation establishment in the developing mouse utricle. (A) In E15.5 utricle, hair bundles are pointing towards each other (arrows) across the line of polarity reversal (LPR, red). The kinocilium is located asymmetrically at the lateral region of the apical HC surface. The MC (yellow), which forms the basal body of the kinocilium, is located more centrally relatively to the DC (orange). Emx2-positive domain is in green. (B) Schematics showing hair bundle orientation in *Emx2* knockout and Gain-of-Function utricles. (C) Model showing asymmetric hair bundle establishment requires the LGN/Inscuteable/Gαi complex. Orientations: A, anterior; M, medial.

Largely based on scanning electron microscopy and immunostaining results, it is thought that the hair bundle is established by first docking of the mother centriole (MC) to the apical center of a nascent HC, where the MC forms the base of the kinocilium (Lu and Sipe, 2016). Then, the kinocilium is relocated from the apical center of the HC to the periphery (Fig. 1C). After the kinocilium acquires its final position, the stereocilia staircase is gradually built next to the kinocilium (Lu and Sipe, 2016, Tarchini et al., 2013, Cotanche and Corwin, 1991, Tilney et al., 1992). The role of the daughter centriole (DC), the inherent partner of the MC, during hair bundle establishment is not known but it is located slightly more peripheral and basal to the MC in mature HCs (Fig. 1A; (Sipe and Lu, 2011)).

Emx2 has a conserved role in reversing hair bundle orientation in HCs of mice and zebrafish (Jiang et al., 2017). However, the timing of Emx2 required to mediate hair bundle reversal is not clear. In other tissues such as the brain, olfactory epithelium and urogenital system, *Emx2* is suggested to function as a patterning gene since the lack of *Emx2* affects regional formation of these tissues (Miyamoto et al., 1997, Pellegrini et al., 1996). Thus, Emx2 could have a similar role in patterning the lateral utricle or specifying the fate of HCs, which indirectly leads to hair bundle reversal. However, Emx2 is known to require the LGN/Insc/Gαi complex in mediating hair bundle reversal (Jiang et al., 2017). The LGN/Insc/Gαi complex forms an asymmetrical crescent on the apical surface of cochlear HCs and this complex is important for guiding the kinocilium to its proper location for cochlear hair bundle establishment and for subsequent stereocilia staircase formation (Fig. 1C; (Ezan et al., 2013, Tarchini et al., 2013, Tarchini et al., 2016)). Therefore, regardless of the mechanisms or timing, Emx2 executes hair bundle reversal by guiding centriole positioning.

In this study, we investigated the timing of Emx2 in hair bundle reversal. We first live-imaged GFP-labeled centrioles in nascent utricular HCs to track the process of hair bundle polarity acquisition. Then, we compared centriole migration trajectories between the Emx2-positive HCs in the lateral and Emx2-negative HCs in the medial utricle to determine whether there is a fundamental difference in their hair bundle polarity establishment (Fig. 2B). We found that there were no obvious differences between medial and lateral HCs in the centriole migration pattern, other than their opposite direction of trajectory. These results indicate that Emx2 has pre-patterned the HCs prior to centriole migration. However, ectopic *Emx2* in naïve medial utricular HCs demonstrated that Emx2 can alter preset centriole trajectories within 12 hours (hrs) and there is a critical time window for centrioles to respond to Emx2. Furthermore, our live-imaging results showed dynamic relationships between the MC and DC, suggesting that the DC may have an active role in guiding the MC. Disruption of microtubule experiments indicate that both centrioles are actively being pulled to its peripheral location via the microtubule network, and ninein, a centrosomal protein, may anchor microtubules to facilitate centriole migration.

**Figure 2.**
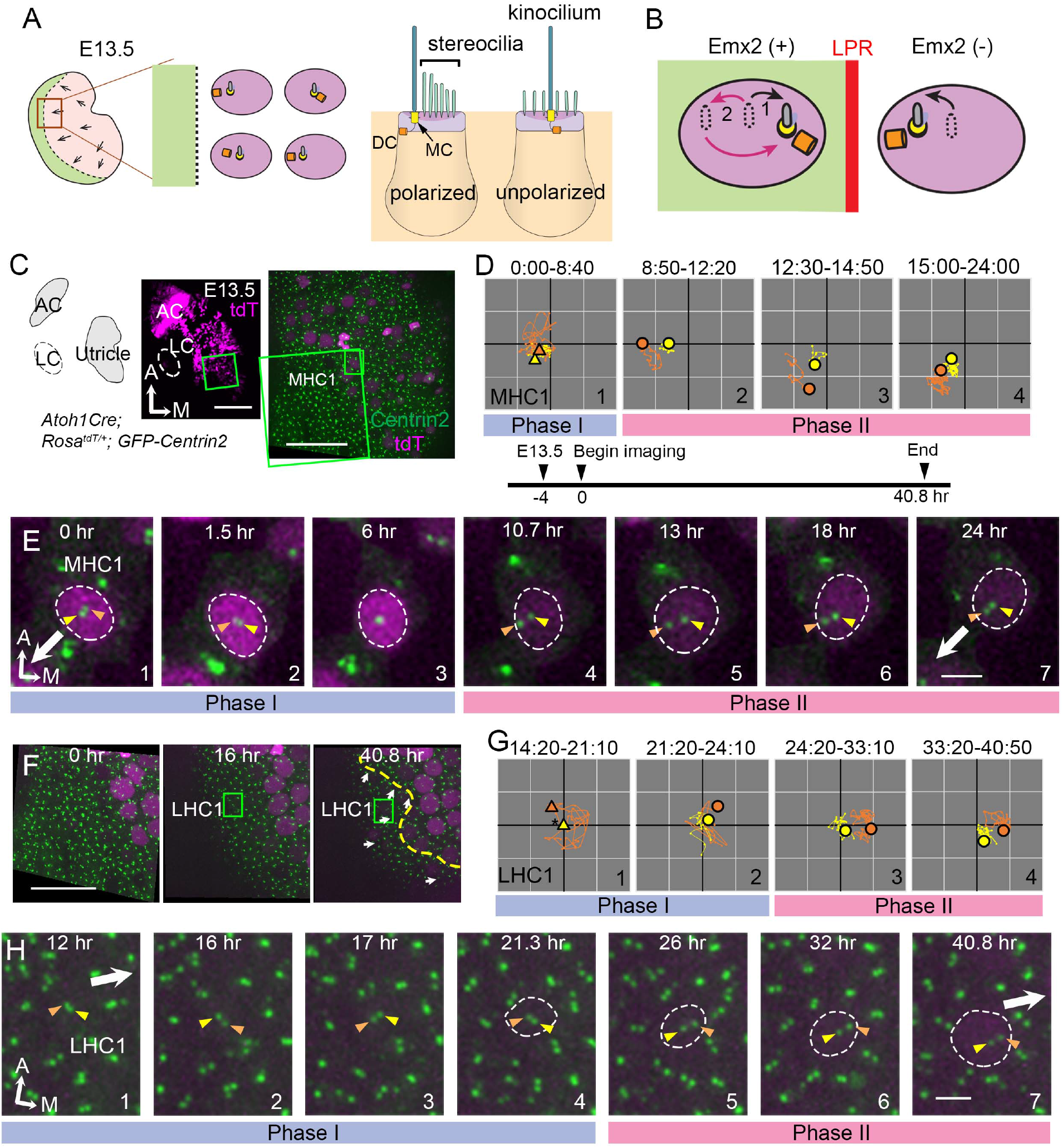
Live imaging of hair bundle establishment in MHC and LHCs based on centriole movements. (A) Utricle schematic at E13.5. The LPR (dotted line) is not apparent since HCs are mostly absent in the Emx2-positive lateral region (green). In the medial utricle, while some HCs are polarized showing the kinocilium located asymmetrically at the lateral region, others are immature and unpolarized with the kinocilium located at the center. (B) In LHCs, MC/kinocilium could migrate directly towards the medial side (black arrow, #1) or it could first migrate toward the lateral side before reversing its direction to the medial side (red arrows, #2). (C) Schematic drawing and images of an *Atoh1^Cre^*; *Rosa^tdT/+^*; *GFP-Centrin2* utricular explant at E13.5 showing locations of MHC1 (small rectangle). (D, E) Centriole trajectories (D) and selected time frames of apical views (E) of MC (yellow) and DC (orange) in MHC1 from time-lapse recording. (D) Yellow and orange triangles represent the beginning positions of respective MC and DC in each time period, and the circled dots represent the final positions in each time period. The yellow and orange lines represent trajectories of the respective MC and DC and are plotted relative to the center of the HC, which is represented by the centroid of the graph. Each small grid is 1.25 × 1.25 μm. All subsequent live-imaging graphs are organized in a similar manner. Initially, the DC is moving vigorously around the MC (Phase I), then the DC starts to move towards the periphery, which is followed by the MC (Phase II). (E) In Phase I (blue bar), the basal body/MC (yellow arrowhead) is located at the center of the HC, whereas the DC (orange arrowhead) moves around the MC. In Phase II (pink bar), the DC starts to migrate towards the lateral periphery of the HC, where the hair bundles should be established in this region of the utricle (white arrow). This migration is followed by the MC. (F-H) LHC1. (F) Selected time frames of the large rectangle area in (C) at the beginning (0:00), middle (16:00) and final frame (40:50) of the recording showing the appearance of LHC1. (G, H) Centriole trajectories (F) and selected time frames of the MC and DC (H) in LHC1. (G,H) tdTomato signal in LHC1 not detectable until 21 hrs into the recording (F, H#1-3) made it difficult to identify the center of the HC. Therefore the position of the MC (yellow triangle) was used as a proxy for the center of the HC (asterisk) for #1 in (G) until the center of HC can be determined in #2-4 in (G). LHC1 shows similar centriole movements as MHC1. Abbreviations: AC, anterior crista; LC, lateral crista; MHC, medial utricular HC; LHC, lateral utricular HC. Scale bars: 100 μm (low mag) and 30 μm (high mag) in (C), 30 μm in (F), 3 μm in (E) and (H).

## RESULTS

### Migration of DC precedes MC in medial utricular HCs during hair bundle establishment

To address how hair bundles are established across the LPR in the macular organs, we first live-imaged centriole movements as a proxy for hair bundle orientation establishment in medial utricular HCs (MHCs) of *Atoh1^Cre^*; *Rosa^tdT/+^*; *GFP-Centrin2* mice at embryonic day (E) 13.5, in which all centrioles are GFP-positive and nascent HCs are tdTomato-positive (Fig. 2A,C-E, 3A-C). At this stage, most of the nascent HCs are located in the Emx2-negative, medial region of the utricle and few are in the Emx2-positive, lateral region (Fig. 2A; (Jiang et al., 2017, Yang et al., 2017)). Some of the MHCs are already polarized with the kinocilium asymmetrically located at the lateral periphery of the apical surface, whereas other immature HCs show the kinocilium at the center of the apical surface (Fig. 2A, S2a). In the course of our experiments, we tracked centriole movements in a total of 32 nascent MHCs for 24 hrs. Figures 2 and 3 illustrate the locations of two of these HCs (Fig. 2C, MHC1, 3A, MHC2) and their centriole movements (Fig. 2D-E, 3B-C). The trajectories of centriole movements (Fig. 2D, 3B) and selected frames of the time-lapse recordings (Fig. 2E, 3C) are shown. The identity of the MC (Fig. 2D-E, 3B-C, yellow color) was determined based on its more apical location within the HC than the DC (Fig. 2D-E, 3B-C, S2a, orange color; (Sipe and Lu, 2011)). The identity of the MC was further validated by its association with the kinocilium marker, Arl13b (Fig. S2a).

**Figure 3.**
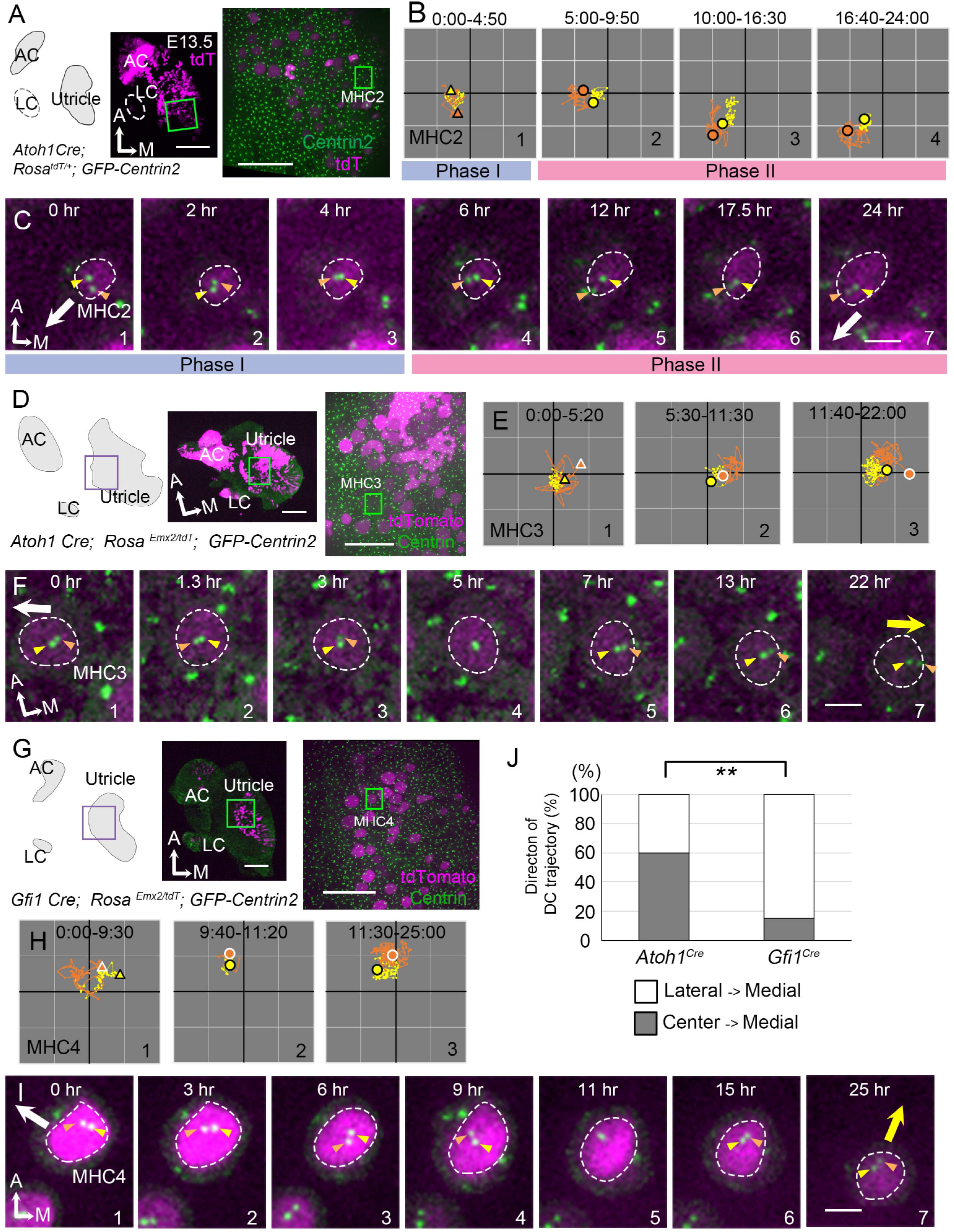
Trajectories of centriole movements in *Emx2* Gain-of-Function MHCs. (A-C) Schematic drawing and images of MHC2 in *Atoh1^Cre^*; *Rosa^tdT/+^*; *GFP-Centrin2* control utricle at E13.5 (A). (B) Trajectory and selected frames (C) from a recording of MC (yellow) and DC (orange) in MHC2. Trajectory is similar to MHC1 in Fig.2. Briefly, the DC is moving vigorously around the MC (Phase I), then the DC starts to move towards the periphery, where hair bundles should be established in this region of the utricle (white arrow). This trajectory is followed by the MC (Phase II). (D-F) Schematic drawing and low and high magnification images of MHC3 in *Atoh1^Cre^*; *Rosa^Emx2/tdT^*; *GFP-Centrin2* utricle at E13.5 (D). (E) Trajectory and selected apical views (F) of MC (yellow) and DC (orange) in MHC3 over-expressing *Emx2*. (E) The DC moves around the MC (#1). Then, the DC starts to move towards the medial side (#2), which is followed by the MC (#3). (F) In #1-4, the DC (orange arrowhead) is moving around the MC (yellow arrowhead) located in the center of the apical HC surface. In #5-7, the DC is moving towards the medial (yellow arrow) instead of the normal lateral (white arrow) utricle. This migration is followed by the MC. (G-I) Schematic drawing, low and high magnification images of MHC4 in *Gfi1^Cre^; Rosa^Emx2/tdT^; GFP-Centrin2* utricular explant (G). (H) Trajectory and selected apical views (I) of the MC and DC in MHC4 over-expressing *Emx2*. Between 0:00-9:30 hr (#1), the centrioles are migrating towards the lateral side of the HC with the DC more lateral than the MC. In #2 (9:40-11:20 hr), the DC starts to change its position to the medial side of the MC, which becomes more apparent in #3 (11:30-25:00 hr). (I) In panels 1-4, the DC (orange arrowhead) is heading towards the lateral side of the utricle (white arrow), then it changes course and moves to the medial side of MC (panels 6-7, yellow arrow). (J) Percentages of DC with two different trajectories in HCs ectopically expressing *Emx2* using either *Atoh1^cre^* or *Gfi^cre^*. Total number of HCs analyzed: *Atoh1^cre^*, n=15; *Gfi^cre^*, n=39. Scale bars: 100 μm (low mag) and 30 μm (high mag) in (A) and apply to (D) and (G), 3 μm in (C) and applies to (F) and (I). ** P<0.01.

Based on tracking and analyses of trajectories and relationships between the MC and DC in MHCs, two phases of centriole movements emerged (Fig. 2D-E, S2b, video 1). In Phase I, the MC (yellow arrowhead) was positioned near the center of HC’s apical surface, whereas the DC (orange arrowhead) moved rapidly and sporadically around the MC (Fig. 2D#1, 2E#1-3, 3B#1, 3C#1-3). The distance between the two centrioles was variable during Phase I (Fig. S2bA), in which the two centrioles could be far apart (Fig. 3C#1) or longitudinally aligned (Fig. 2E#3, 3C#3). The speed of the DC was also faster, up to 1.2 μm per a 10-minute time frame (orange), whereas the speed of the MC (yellow) was more stable, not exceeding 0.5 μm per time frame after correcting for HC drift (Fig. S2bB, MHC1: p=1.1×10^−6^, MHC2: p=0.015). By contrast, Phase II was characterized by the DC showing directional movements towards the lateral utricle (Fig. 2D#2, 2E#4-5, 3B#2, 3C#4-5) where the kinocilium of MHCs will subsequently reside in the lateral periphery (white arrow). Then, the MC migrated towards the direction of the DC (Fig. 2D#3-4, 2E#6-7, 3B#3-4, 3C#6-7). The DC continued to move faster than the MC in Phase II (Fig. S2bB, MHC1: p=2.3×10^−5^, MHC2: p=4.7×10^−6^) and the average distance between the two centrioles in Phase II was significantly further apart than in Phase I (Fig. S2bA, MHC1: p=8.0×10^−8^, MHC2: p=2.0×10^−5^).

### Lateral HCs show trajectories of MC and DC similar to medial HCs

Hair bundles in the lateral utricle are in opposite orientation from the default hair bundles in the medial utricle (Fig. 1A). We investigated whether centrioles in the Emx2-positive HCs migrate directly to their destinated position in the medial periphery or they first migrate to the lateral position before relocating to the medial destinated position (Fig. 2B). Since most of the HCs in the lateral utricle initiate terminal mitosis at E14.5 or later (Jiang et al., 2017), live-imaging of E13.5 utricular explants was extended to 41 hrs (Fig. 2F-H, S2cA-E). At the beginning of recordings, tdTomato-positive HCs were only found in the medial utricle. Twenty hrs into imaging, several HCs emerged in the lateral region of the utricle and their centrioles were positioned in the medial periphery of the apical surface of the HCs by the end of the recording (Fig. 2F, LHC1, S2cA, LHC2). Based on the positions of HCs at the end of the recordings, we re-traced and analyzed the centriole trajectories of 9 lateral HCs (LHCs). Trajectories of centrioles in LHCs can also be grouped into two phases similar to MHCs. In Phase I, the DC moved sporadically around the MC before or soon after the tdTomato signal was evident (Fig. 2G#1-2, 2H#1-4, S2cB#1-2, S2cC#1-4, video 2). Then, in Phase II, the DC started to migrate towards the medial side, which was followed by the MC (Fig. 2G#3-4, 2H#5-7, S2cB#3-4, S2cC#5-7). Similar to MHCs, the average distance between the two centrioles was variable but closer in Phase I than in Phase II (Fig. S2cD, LHC1: p=2.7×10^−11^, LHC2: p=1.7×10^−17^). Average moving speed of the DC was faster than that of the MC in each phase, although three out of nine LHCs failed to show a significant difference in speed between the two centrioles in Phase II (Fig S2cE, LHC1: Phase I p=0.0078, Phase II p=0.028, LHC2: Phase I p=0.048, Phase II p=0.19).

Thus far, our analyses revealed that MHCs and LHCs show similar centriole trajectories in reaching their destinations in opposite sides of the HC. The two centrioles in the LHC moved directly toward the medial periphery (Fig. 2B#1) without first reaching the lateral periphery like in the MHC and then relocate to the medial periphery (#2). These results indicate that Emx2 has already exerted its effects on the LHCs prior to hair bundle establishment. Notably, *Emx2* transcripts are detected in the lateral region at E11.5, three days ahead of HC formation that begins at E14.5 (Fig. S2d; (Jiang et al., 2017)). Taken together, these results suggest that the mechanism of Emx2 in altering hair bundle orientation in the lateral utricle could be indirect via possible regional patterning and/or HC fate determination.

### Reversing hair bundle orientation by ectopic *Emx2*

To gain further insight into the mechanism of Emx2 in altering hair bundle orientation, we investigated the time it takes for ectopic Emx2 to reverse hair bundle orientation in MHCs (Jiang et al., 2017). Since *Emx2* transcripts are detected in the lateral utricle three days earlier than LHC formation (Fig S2d; (Jiang et al., 2017)), we reasoned that if endogenous Emx2 mediates hair bundle reversal via patterning or cell-fate change, effects of ectopic *Emx2* may be similar and should take days to reverse hair bundle orientation. We crossed two different strains of *cre* mice, *Atoh1^Cre^* or *Gfi1^Cre^* to *Rosa^Emx2^* mice, which resulted in some offspring showing specific expression of *Emx2* in all the HCs. Both Atoh1 and Gfi1 are the transcription factors important for HC formation (Bermingham et al., 1999, Wallis et al., 2003), and lack of *Atoh1* or *Gfi1* results in loss of HCs. Atoh1 is the earliest known transcription factor that commits HC fate in the inner ear (Bermingham et al., 1999, Zheng and Gao, 2000). However, *Atoh1* expression is not affected in *Gfi1* knockout mice, suggesting that Gfi1 is required later than Atoh1 during HC differentiation (Wallis et al., 2003). Thus, the induction of *Emx2* using *Gfi1^Cre^* is expected to be later than that of *Atoh1^Cre^* in the HC lineage.

We first live-imaged *Atoh1^Cre^; Rosa ^Emx2/tdT^; GFP-Centrin2* utricles (Fig. 3D-F), in which all MHCs showed opposite hair bundle orientation from controls by E15.5 (Fig. S3a). Live-imaging results showed that the trajectory of centrioles in MHCs ectopically expressing *Emx2* showed the DC moving around the MC sporadically (Fig. 3E#1, 3F#1-4), similar to normal MHCs at Phase I (Fig. 3B#1, 3C#1-3). Then, the DC migrated toward the medial periphery (Fig. 3E#2-3, 3F#5-7, yellow arrow, video 3), opposite from the normal lateral direction (Fig. 3F, white arrow) and controls (Fig. 3B-C). This trajectory was followed by the MC. This pattern of centriole migration from center of the HC to the medial edge occurred in 60% of the HCs analyzed (Fig. 3J). The remaining HCs exhibited a pattern that is similar to the *Gfi1^Cre^*; *Rosa^Emx2/tdT^* MHCs described below, suggesting that there are two modes of centriole trajectory.

In *Gfi1^Cre^*; *Rosa^Emx2/tdT^* utricles (Fig. 3G-I), in which all the hair bundle orientation in MHCs are known to be reversed (Jiang et al., 2017), live imaging results showed that some of the centriole trajectories were different from those in *Atoh1^Cre^*; *Rosa^Emx2/tdT^* utricles (Fig. 3G-J, p=0.0010, Fig. S3bA-C). At the beginning of the recording, *Gfi1^Cre^*; *Rosa^Emx2/tdT^* MHCs analyzed already showed tdTomato expression and the DC was asymmetrically located towards the lateral side (Fig. 3H#1, 3I#1, MHC4, S3bC-C”#1, MHC5-7, white arrow, video 3), resembling MHCs at Phase II (Fig. 3C). Within 3 hrs (Fig. S3bC’#1-3, MHC6) to 9 hrs (Fig. 3I#1-4, MHC4, S3bC” #1-4 MHC7) of recordings, the DC remained lateral to the MC. Thereafter, the distance between the DC and MC was reduced and sometimes the two centrioles were transiently aligned longitudinally (Fig. 3I#5 MHC4, S3bC#5 MHC5, C”#5-6 MHC7). Then, the DC switched position to the medial side of MC (Fig. 3H#3, 3I#6-7, S3bC-C”, yellow arrow). These results suggest that in *Gfi1^Cre^*; *Rosa^Emx2/tdT^* MHCs, the centrioles are initially aligned in the normal lateral positions primed for hair bundle establishment but upon activation of *Emx2* transcription mediated by *Gfi1*-driven Cre, the two centrioles reverse their positions to the medial side. This lateral to medial centriole reversal pattern was also observed in *Atoh1^Cre^*; *Rosa^Emx2/tdT^* utricles but at a lower frequency than MHCs of *Gfi1^Cre^*; *Rosa^Emx2/tdT^* (Fig. 3J, 40% vs 84.6%, p=0.0010). These frequency differences in migration patterns between the two *cre* lines are consistent with the notion that *Emx2* activation using *Gfi1^Cre^* is later than *Atoh1^Cre^*, thus a higher percentage of centrioles in *Gfi1^Cre^* samples were observed to migrate from the destinated lateral periphery towards the medial rather than directly from the central towards the lateral position (Fig. 3J, 84.6% vs 15.4%). Furthermore, our results indicate that centriole trajectory during both Phase I and II is plastic and can be altered by Emx2.

### Infections with AAV-Emx2 reverse hair bundle orientation

Our time-lapse recordings showed that the longest time observed for *Gfi1^Cre^*; *Rosa^Emx2/tdT^* MHCs to reverse centriole positions from the lateral to medial periphery was approximately 12 hrs (Fig. 3, MHC4, S3b, MHC7) suggesting that Emx2 could exert its effects on hair bundle reversal within hours. However, the precise time frame between the onset of *Emx2* expression driven by *Gfi1-*Cre and reversal of centriole positions remains unclear. To further investigate the time required by Emx2 to alter hair bundle orientation, we ectopically expressed *Emx2* in utricular explants using AAV2.7m8 adeno-associated virus, which has been shown to infect cochlear HCs efficiently (Isgrig et al., 2019). We infected *GFP-Centrin2* utricular explants at E13.5 with an AAV2.7m8 adeno-associated viral vector, AAV2.7m8-CAG-Emx2-P2A-tdTomato (AAV-Emx2-tdT), in which both *Emx2* and *tdTomato* transcripts were driven under the universal CAG promoter. Forty-eight hrs after infection, approximately 70% of total HCs were infected (Fig 4C, 71.1 ± 6.3 % with AAV-Emx2-tdT), which was similar to AAV-tdT controls (72.8 ± 5.6 %). The void of anti-β2-spectrin staining indicates the kinocilium position in the apical HC surface and reveals the hair bundle orientation (Deans et al., 2007). In AAV-tdT controls, infected HCs show normal hair bundle orientation (Fig. 4A,C). However, 13.9 ± 7.2 % of HCs infected with AAV-Emx2-tdT showed opposite kinocilium position (Fig 4B,C) versus none in controls, indicating that AAV-Emx2-tdT is sufficient to reverse hair bundle orientation.

**Figure 4.**
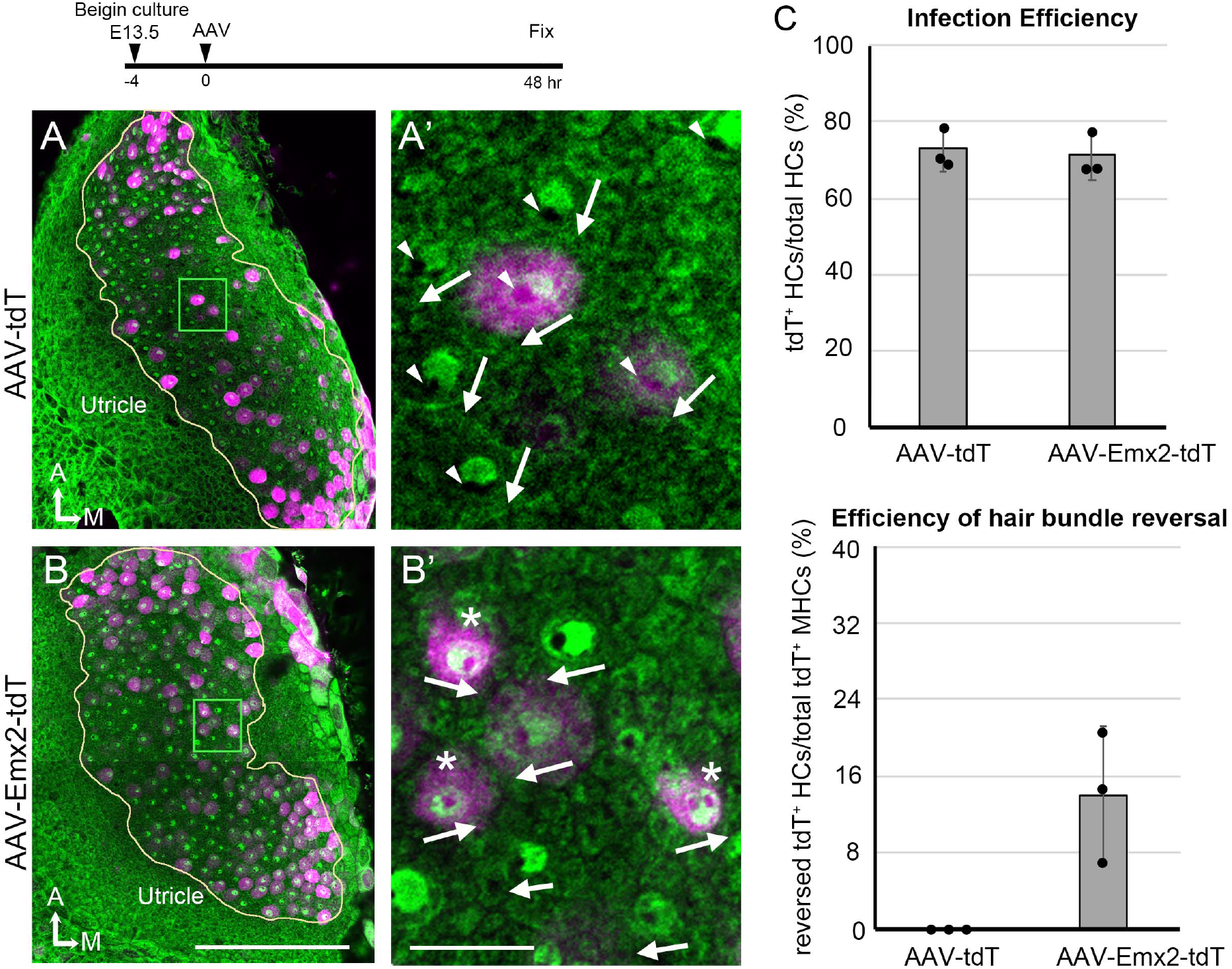
AAV-Emx2 infections reverse hair bundles in MHCs. (A, A’) Low (A) and high magnification (A’) images of the rectangular area of a control explant infected with AAV-tdT (tdT: magenta) and stained with anti-β2-spectrin antibodies (green), in which the absence of staining indicates the kinocilium location (arrowhead) and HC orientation (arrows). The two AAV-tdT infected HCs (magenta) show hair bundle orientation similar to non-infected HCs. (B,B’) Low (B) and high magnification (B’) images of the rectangular area of an utricular explant infected with AAV-Emx2-tdT (tdT: magenta), stained with anti-β2-spectrin antibodies (green). Some of the AAV-Emx2-tdT infected HCs show reversed hair bundle orientation (asterisk) from the rest of the non-infected HCs. (C) Efficiency of viral infections and efficiency of hair bundle reversal among infected HCs (n=3 experiments for each condition). Error bars represent SD. Scale bars: 100 μm in (B) and applies to (A), and 10 μm in (B’) and applies to (A’).

Next, we live-imaged the centriole reversal process in AAV-Emx2-tdT infected *GFP-Centrin2* utricular cultures (Fig. 5A). At 24 hrs after viral infection, the majority of the cells in the utricular explants were tdTomato-negative (Fig 5B) but many HCs turned on tdTomato by 36 hrs after infection (Fig. 5E,F). At the beginning of recordings, most of the infected MHCs showed the DC lateral to the MC, even though the outline of the HCs was not evident yet due to the lack of tdTomato signal (Fig. 5C-F#1, video 4). As tdTomato expression became apparent, the peripheral location of the two centrioles was confirmed, indicating that these cells were at the end of Phase II (Fig. 5E-F, MHC8, 21hrs, MHC9, 13 hrs). Then, the two centrioles moved to align longitudinally with each other transiently (Fig. 5E-F, small panel insets, 22.3 hrs for MHC8 and 15 hrs for MHC9), followed by the DC moving medial to the MC (Fig. 5C#3, 5E 23.3-25.3 hrs, MHC8, 5D#3, 5F 19.5-25.3 hrs, MHC9), suggesting a change in the course of DC trajectory from lateral to medial periphery. Quantification of tdTomato signal that was above background in infected cells showed that tdTomato signals were elevated by 9-12 hrs of imaging (Fig. 5A,G, 1.5 days after infection), although the signals may not be apparent in the single time frame images (Fig. 5E,F). Then, DC positional reversal occurred within 10-12 hrs after the detectable tdTomato signals. Using positive tdTomato signals as a proxy for Emx2 activation, these results suggest that Emx2 can reverse centriole trajectory within 10-12 hrs. Thus, both the viral approach using AAV and genetic approach using *Gfi1^Cre^*; *Rosa^Emx2/tdT^* utricles suggest that Emx2 is able to mediate hair bundle reversal within a short period of time (Fig. 5H) despite onset of endogenous *Emx2* expression occurring three days prior to HC formation (Fig. S2d).

**Figure 5.**
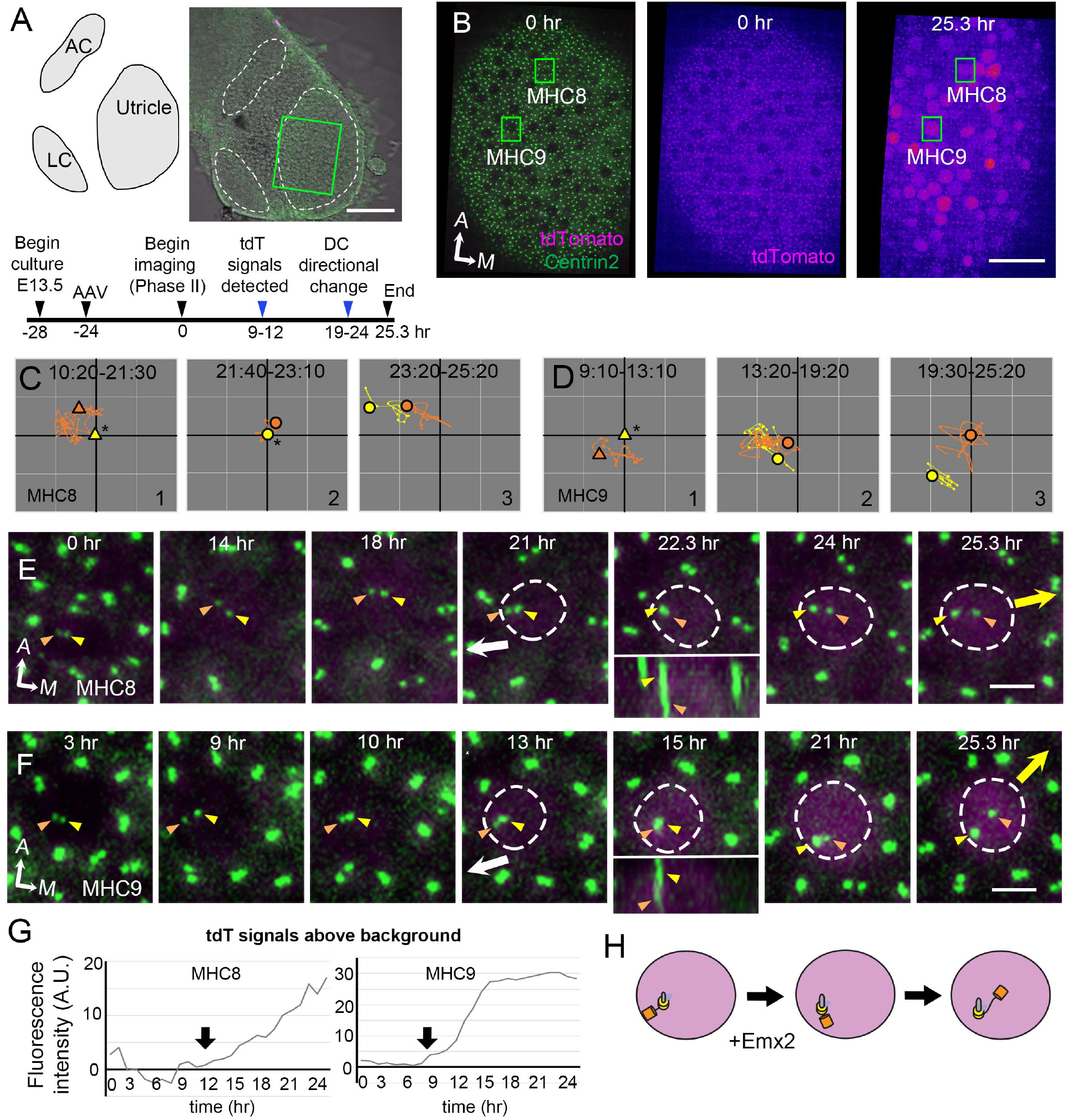
Altered DC trajectory in MHCs infected with *AAV-Emx2*. (A) Schematic and low magnification image of the *GFP-Centrin2* utricular explant infected with AAV-Emx2-tdT at E13.5. The timeline of experimental treatments (black arrowheads) and observations (blue arrowheads) are shown. (B) A utricular explant at the beginning (0:00) and end (25:20) of a time-lapse recording showing clear tdTomato-positive cells by the end of recording. (C, D) The trajectory of the MC and DC in MHC8 and MHC9. Initially, the DC (orange triangle, #1) is positioned lateral to the MC (yellow triangle) at the center (asterisk). Then, the DC (orange dot) moves sporadically around MC (yellow dot, #2), followed by DC moving medial to the MC in #3. (E, F) Selected frames of the recording of MHC8 and MHC9. At the beginning, the DC is located by the lateral side(white arrow) of each HC. Then, the DC overlaps with the MC briefly at 22.3 hr for MHC8 and 15 hr for MHC9 during recording (insets showing side views), followed by the DC moving medial to the MC towards the medial direction (yellow arrow). (G) tdTomato expression compared to the background level, indicating tdTomato signals exceeded background after 12 (MHC8) and 9 hrs (MHC9) of recordings (arrows). (H) Schematic of centriole movements in the presence of Emx2. Scale bars: 100 μm in (A), 30 μm in (B) and 3 μm in (E) and (F).

### Microtubules are required to stabilize the asymmetrical location of the centrioles

Thus far, our live-imaging results indicate that the migration of the DC always preceded that of the MC under either control or treated conditions. To identify potential qualitative differences between the two centrioles that could account for the migration pattern, we investigated their association with microtubules and proteins related to microtubule nucleation and anchoring. Using SiR-tubulin, a cell permeable fluorogenic probe for microtubules, we labeled microtubules in live *Atoh1^Cre^*; *Rosa^tdT/+^*; *GFP-Centrin2* utricular cultures. We showed that both centrioles were associated with microtubules in tdTomato-positive HCs determined to be Phase II, based on the position of the centrioles (Fig. 6A). Immunostaining with anti-γ-tubulin antibodies indicated that both centrioles in Phase II HCs are associated with microtubule nucleation (Fig. 6B). Ninein is a centrosomal protein that has both microtubule nucleation and anchoring functions (Delgehyr et al., 2005) and it is preferentially associated with the MC than the DC in somatic cells, serving its microtubule nucleation role (Betleja et al., 2018, Piel et al., 2000). In Phase I HCs, in which centrioles are centrally located, ninein staining was concentrated by the two centrioles (Fig. 6C, Phase I, n=6). As DC started to migrate towards the periphery and the two centrioles became further apart, ninein staining became broadly distributed surrounding both centrioles (early Phase II, n=17). However, ninein staining was concentrated at the centrioles again by the end of Phase II (Fig. 6C, n=21). This broad distribution of ninein staining beyond the centrioles during centriole migration suggests that ninein may serve as microtubule anchoring during this period.

**Figure 6.**
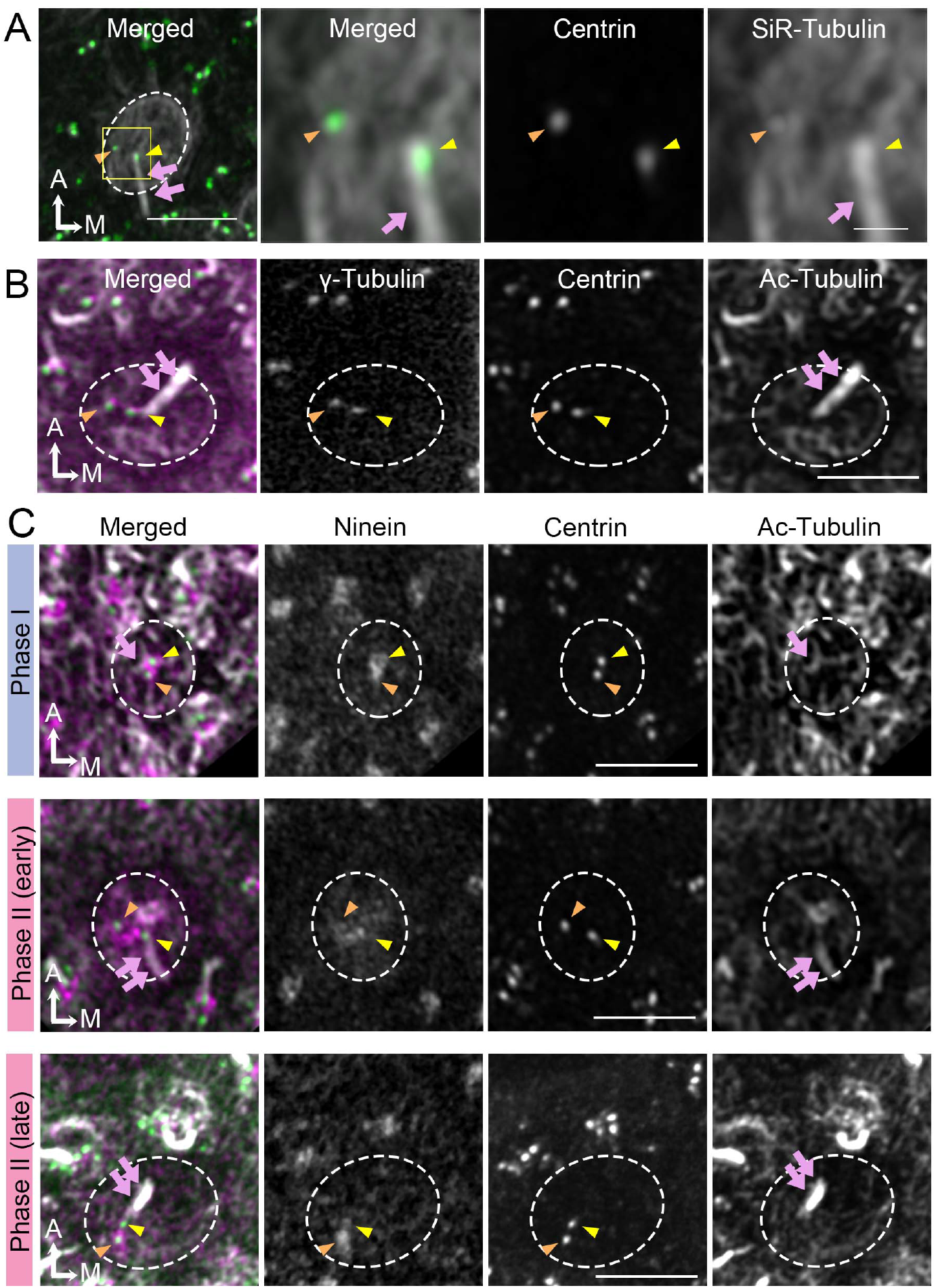
Broaden ninein localization during centriole migration. (A) SiR-tubulin (white) labeling of an MHC of E13.5 *Atoh1^Cre^*; *Rosa^tdT/+^*; *GFP-Centrin2* utricle at Phase II. The magnified views of the yellow rectangular area in the left panel are shown in the three right panels, illustrating that both centrioles (green in the merged picture) are associated with the microtubule network. MC, yellow arrowhead; DC, orange arrowhead; kinocilium, pink arrows. Dotted white lines indicate apical surface of the HC. (B) γ-tubulin expression of an MHC at Phase II. DC and MC show similar expression of γ-tubulin (magenta in the merged picture). Acetylated tubulin labels mature microtubule and the kinocilium. (C) Immunostaining of ninein in MHCs in Phase I, early and late Phase II. Phase I (outlined) and late Phase II HCs show centrosomal ninein (magenta on merged picture) staining. At the beginning of Phase II, ninein staining is diffuse and broader than centrioles. Scale bars: 3 μm in the left panel in (A) and applies to (B-C’), 1 μm in the three right panels of (A).

Previous reports in cochlear explants proposed that microtubule plus ends attached to the LGN/Insc/Gαi complex pulls the MC/kinocilium through microtubule shortening and/or dynein mediated mechanism to the periphery (Ezan et al., 2013). Blocking Gαi with pertussis toxin disrupted the microtubule plus-end binding protein, EB1 and the kinocilium positioning. We tested this hypothesis in our live utricular culture and asked whether the DC behaves in a similar manner as MC. We treated utricular explants with nocodazole, which disrupts the microtubules by binding to free tubulin dimers and inhibits microtubule polymerization (Hoebeke et al., 1976). We focused our analysis on MHCs that were at the end of Phase II, which showed both centrioles located peripherally in the lateral region (Fig. 7A-C). Nocodazole was introduced at 1.5 hrs into live-imaging. Shortly after the addition of nocodazole, positions for both centrioles were affected and they returned to the center of the HC with the DC traveling more rapidly and sporadically around the relatively stable MC (Fig. 7B,C, 1.7-2.5 hrs), resembling the behavior observed in Phase I of nascent HCs (Fig. 2). Once nocodazole was washed out, two centrioles returned to the lateral periphery within a few hrs with the DC moving ahead of the MC (Fig. 7C, 2.5-8 hrs, video 5). The acetylated-tubulin staining of HCs, which labels stable microtubules, showed a loss of tubulin arrays along with mislocalized centrioles from the periphery during nocodazole treatment (Fig. 7D). After nocodazole removal, tubulin arrays were re-established and centriole positions in the periphery recovered (Fig 7E). These results indicate that the maintenance of both the MC and DC in the periphery is dependent on an intact microtubule network and an active force that pulls both centrioles to the periphery.

**Figure 7.**
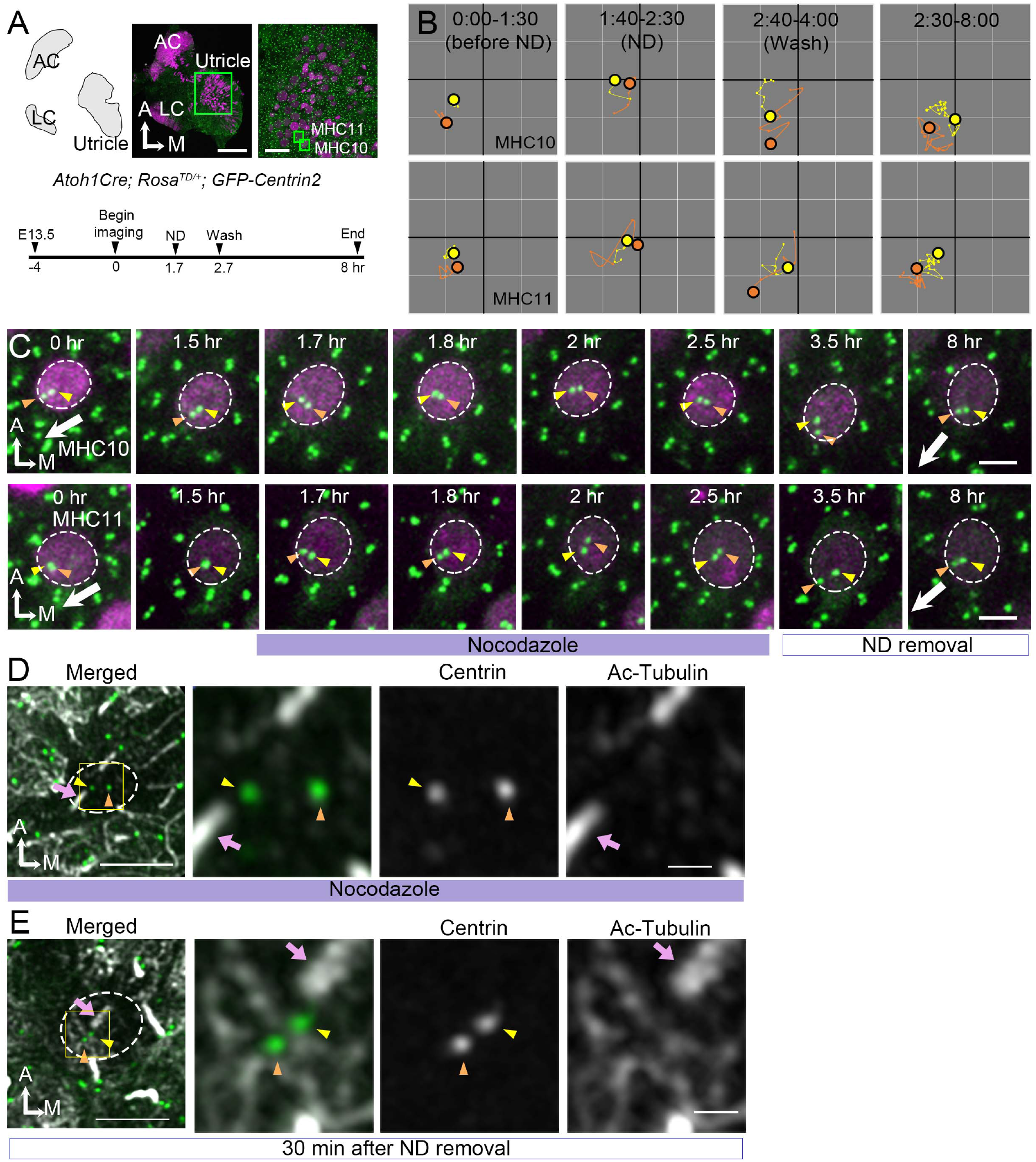
Centrioles lose their lateral position after nocodazole treatments. (A) Schematic, low and high magnifications of an *Atoh1^Cre^*; *Rosa^tdT/+^*; *GFP-Centrin2* utricular explant. The timeline indicates the experimental treatments. (B) The trajectory of MC and DC in MHC10 and MHC11. Before the introduction of nacodazole (ND, 0:00-1:30 hr), the DC is located more peripherally than the MC, towards the lateral edge of HC (Phase II). The position of the DC and MC changes during ND treatment and they move back to the center of the HC with DC moving ahead of MC. After removal of the ND, centrioles return to the polarized location again with DC moving ahead of MC. (C) Selected frames of MHC10 and MHC11 before (0 hr), during (1.7-2.5 hr) and after washing out (3.5-8 hr) of ND. Before ND treatment, the DC is positioned lateral to the MC towards the direction where hair bundles will be established (white arrow). This relationship is disrupted during ND treatment with both DC and MC relocating to the center of the HC and the DC is more medial than the MC at the end of drug treatment (2.5 hr). After drug removal, the DC returns to its original position within 1 hr, which is followed by the MC (8hr). (D) The nocodazole-treated MHC shows mispositioned centrioles (MC, yellow arrowhead; DC, orange arrowhead) that are no longer asymmetrically located in the periphery. The three panels on the right are merged, single centrin and acetylated tubulin images of the rectangular area in the left panel. Tubulin arrays are absent in the cytoplasm of HC. (E) An MHC after removal of ND for 30 min shows centrioles returning to their peripheral location. The three panels on the right are magnifications of the rectangular area in the left panel. Tubulin arrays (white) are radiating from the centrioles to the periphery. Scale bars: 100 μm (low mag) and 30 μm (high mag) in (A), 3 μm in (C) and the first panel in (D) and (E), and 1 μm in high mag of (D) and (E).

## Discussion

### Hair bundle establishment in HCs

Our live-imaging study is the first extensive time-lapse imaging of hair bundle acquisition in mammalian HCs. Imaging results of centriole migration in utricular HCs are consistent with a previous model extrapolated from results of SEM that the kinocilium starts out in the center of a HC before reaching its peripheral destination for hair bundle establishment (Dabdoub et al., 2003, Lu and Sipe, 2016, Denman-Johnson and Forge, 1999). Based on results in the chicken basilar papilla, it was proposed that the kinocilium undergoes fairly extensive migration along the periphery of the HC before reaching its final destination (Fig. 1A; (Cotanche and Corwin, 1991, Tilney et al., 1992). However, more recent studies in the mouse cochlea suggest that the MC/kinocilium takes a more direct route from the center of the HC to its final destination in the periphery (Dabdoub et al., 2003, Lu and Sipe, 2016, Denman-Johnson and Forge, 1999, Montcouquiol et al., 2003). Our results are consistent with findings in the mouse that the MC/kinocilium and DC take a direct path to their destination in the periphery. Additionally, we found an intriguing centriole migration pattern for hair bundle establishment as discussed below.

It has been proposed that the migration of kinocilium to its destinated location is achieved through an active force on centrioles via the microtubule network. This active force is suggested to be exerted through LGN/Insc/Gαi complex on microtubules by recruiting Lis1/dynein and/or shortening of microtubule (Ezan et al., 2013, Lu and Sipe, 2016, Tarchini and Lu, 2019). Our nocodazole results supported this active force hypothesis. Furthermore, disruption of the microtubule network caused the centrioles to promptly return to their initial central location rather than to remain stationary at the periphery of HCs suggests that Lis1/dynein is a more likely mechanism than microtubule shortening. We reasoned that disruption of microtubule shortening should leave centrioles in position rather than return to its original central position of the HC. Furthermore, our nocodazole results indicate that the DC is likely to be under similar control of the LGN/Ins/Gαi complex as the MC.

The directed migration of the centrioles depends on oriented microtubule arrangement of the minus ends at the centrioles and the plus ends at the periphery. This arrangement raises the possibility that proper anchoring of microtubules within HCs is an important factor. Among the anchoring proteins, we focused on ninein, which has both microtubule nucleation and anchoring functions (Delgehyr et al., 2005). It functions as a microtubule nucleation by docking γ-tubulin to the centrosomes and it also anchors microtubules in non-centrosomal site. For example, in pillar cells (supporting cells) of the cochlea, ninein has a non-centrosomal location in adherens junctions, which serve as microtubule anchors (Mogensen et al., 2000, Moss et al., 2007). Notably, ninein distribution was no longer concentrated at the centrioles and became broader during centriole migration. This broaden ninein distribution during centriole migration suggests that microtubule organization is different during centriole migration and raises the possibility that ninein may function as microtubule anchors in addition to microtubule nucleation during centriole migration (Fig. 8).

**Fig. 8.**
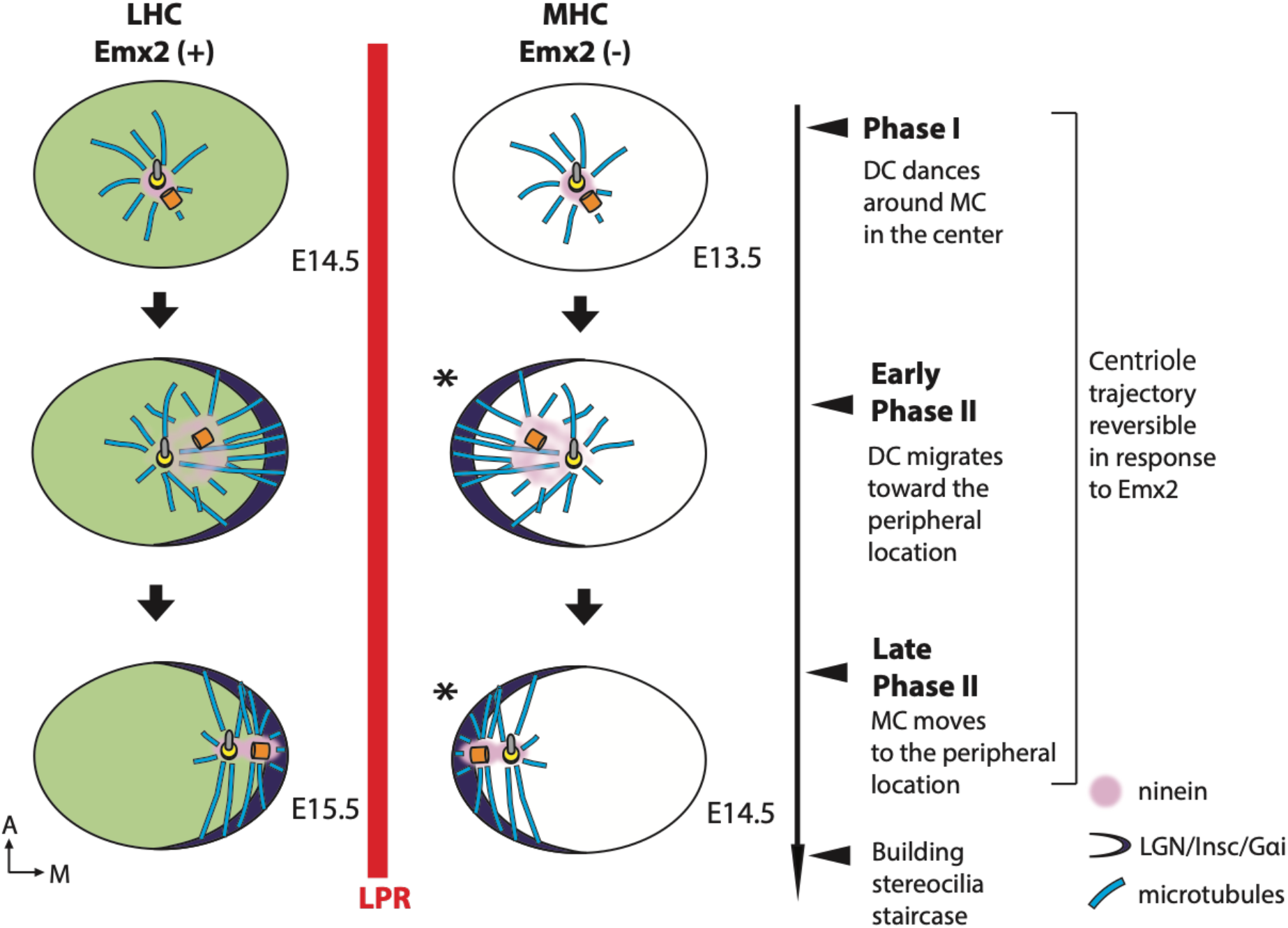
Summary of centriole trajectory during hair bundle establishment in nascent utricular HCs. In Emx2-negative MHCs, Phase I is represented by MC located in the apical center of HCs with the DC dancing around the MC. Ninein is associated with the centrioles, which serves as the nucleation center for microtubules. In early Phase II, when DC migrates toward the peripheral side, the MC starts to follow the direction of the DC and the peripheral crescent LGN/Insc/Gαi complex starts to be established (Ezan et al., 2013, Tarchini et al., 2013). The broad distribution of ninein surrounding both centrioles is likely to anchor the minus end of microtubules and facilitates centriole migration to the periphery. By the end of Phase II, both MC and DC are located in the periphery and ninein becomes restricted to the centrioles again. Centriole trajectory in both Phase I and II are reversible in the presence of Emx2 but responsiveness to Emx2 decreases over time (Jiang et al., 2017). Emx2-positive LHCs (green) show similar but opposite trajectory pattern of centriolar migration. Nevertheless, LGN/Insc/Gαi may be dispensable for hair bundle establishment in the MHCs since blocking Gαi with pertussis toxin does not appear to affect bundle orientation (asterisk).

### Relationship between DC and MC migration

Many well described morphological and functional features distinguish between the MC and DC (Pelletier and Yamashita, 2012, Fujita et al., 2016). In addition to the function of DC maturing into an MC during the cell cycle, the function of the DC in differentiating cells is only beginning to be understood (Loukil et al., 2017, Betleja et al., 2018, Gottardo et al., 2015). Other than the DC being actively inhibited to form the cilium, recent results indicate that the proximity of the DC to the MC is also important for primary cilium formation (Loukil et al., 2017). Here, in nascent HCs, we show that when the MC was located at the center during Phase I, the DC was observed to move sporadically around the relatively stationary MC (Fig. 8). This phenomenon has been described in several vertebrate somatic cell lines (Piel et al., 2000). In these cell lines, the MC, which is associated with a microtubule network is stationary, whereas the DC, though associated with the microtubule nucleation marker, γ-tubulin, is more mobile. The functional significance of the mobile DC, however, remains unclear except that this behavior is regulated by the cell cycle and attenuates as cells transition from G1 to S phase.

During Phase II of the centriole migration, the DC invariably moved ahead of the MC to reach the peripheral destination. This pattern of the DC preceding the MC in migration was observed repeatedly in HCs under all conditions investigated such as during normal centriole migration, nocodazole treatments and recovery, and ectopic *Emx2* activation, suggesting that the migration of the DC is related to the MC. Although little is known about the relationships between the DC and MC in HCs, an inner ear conditional knockout of *Kif3a*, which encodes an intraflagellar transport protein, shows misplaced location and relationship between the MC and DC in cochlear HCs (Sipe and Lu, 2011).

Several scenarios could account for the observed behavior of the DC moving ahead of the MC in HCs. One possibility is that the two centrioles move independently of each other. Since each centriole is associated with a microtubule nucleation center as indicated by their association with γ-tubulin staining (Fig. 6B), each can be independently pulled by the microtubule-dynein system to their destinations in the peripheral cortex. The faster and higher mobility of the DC may simply be due to the MC being restricted by the attached cilium (Paintrand et al., 1992). An alternative scenario is that the MC is being dragged to the periphery by the DC via the intercentrosomal linkers between MC and DC, which are made of rootletin filaments (Yang et al., 2006). The sporadic movements of the DC around the MC in Phase I could also be regulated by the intercentrosomal linkers. In other systems, the length of these linkers can change and disintegrate based on maturation of the centrioles during the cell cycle (Bahmanyar et al., 2008, Mardin et al., 2010). However, little is known about the regulation and possible functions of these linkers in post-mitotic cells including HCs. Based on our findings, we speculate that the DC has an active role in guiding the MC/kinocilium to its proper location in differentiating HCs, in addition to its role in regulating ciliogenesis.

### The role of Emx2 in reversing hair bundle orientation

In zebrafish lateral line, Emx2 regulates neuronal selectivity as well as hair bundle orientation (Ji et al., 2018). In the mouse utricle, onset of *Emx2* expression is well ahead of the emergence of HCs (Fig. S2d). Therefore, Emx2 may have a role in regional patterning and/or HC fate specification that indirectly lead to hair bundle reversal. Our live-imaging results demonstrating that HCs are already pre-patterned by Emx2 prior to centriole movements supports this hypothesis (Fig. 8). Nevertheless, our ectopic *Emx2* experiments using AAV indicate that tdTomato signal was detectable within 36 hrs of AAV-Emx2 infection (Fig. 5). This timeframe of sequential transcriptional and translational events to yield detectable tdTomato signal is comparable to other mammalian systems. Under the assumption that Emx2 is synthesized in a comparable time frame as tdTomato, hair bundle orientation reversal occurred relatively quickly within approximately 12 hours of detectable tdTomato. These results suggest that while Emx2 may have other functions in the utricle, its bundle reversal effect is likely to be direct and does not require multiple cascades of transcriptional and translational events. Additionally, both the genetic and AAV viral approaches indicate that during these early phases of centriole migration in hair bundle establishment, the system is plastic and responsive to Emx2 (Fig. 8). However, the time-window of centrioles’ responsiveness to Emx2 is critical as ectopic *Emx2* after E15.5 only has a limited effect on hair bundle reversal in naïve HCs (Jiang, Kindt & Wu 2017). This critical time-window may also attributed to the low frequency of hair bundle reversal observed in AAV-Emx2-tdT infected HCs (Fig. 4).

Furthermore, our results showed that the DC may have an active role in guiding the MC to its designated location in the HC periphery and a positive force is required to actively maintain this peripheral centriole positioning. These findings provided insights into the regulation of centriole dynamics during hair bundle establishment.

## Materials and methods

### Mouse

All animal experiments were conducted according to NIH guidelines and under the Animal Care Protocol of NIDCD/NIH (#1212-17). *GFP-Centrin2* mice were obtained from Xiaowei Lu at University of Virginia (PRID:MGI:3793421), *Atoh1-Cre* mice from Bernd Fritzsch at University of Iowa (PRID:MGI: 3775845), and *Gfi1-Cre* mice from Lin Gan at Augusta University (PRID:MGI:4430258; (Yang et al., 2010)). The *Rosa26R^Emx2^* mouse was generated by knocking in the cassette *attb-pCA promoter-lox-stop-lox-Emx2-T2A-Gfp-WPRE-polyA-attb* to the *Rosa* locus described previously (Jiang et al., 2017). *Rosa26R^tdTomato^* were purchased from Jackson laboratory (RRID:IMSR_JAX:007914, (Madisen et al., 2010)). *Atoh1^Cre^*; *Rosa^tdT/+^* control specimens for live imaging was generated by crossing *Atoh1^Cre^*; *Rosa^tdT/tdT^* males with *GFP-Centrin2*^+/−^ females. Emx2 gain-of-function specimens was generated by crossing *Atoh1^Cre^*; *Rosa^tdT/tdT^* or *Gfi1^Cre^*; *Rosa^tdT/tdT^* males with *GFP-Centrin2^+/−^*; *Rosa^Emx2/Emx2^* females. In addition to tdTomato signals, HC identity in the live-images was confirmed based on the round or oval shape of the cuticular plate and its apical position of the nucleus within the epithelium relative to nuclei of the supporting cells.

### Live imaging

The mouse utricle together with anterior and lateral cristae for orientation were dissected from E13.5 mouse inner ears. The harvested tissue was mounted on Cell-Tak (Corning, NY, NY)-coated coverslips (Belyantseva, 2016) in DMEM/F12 (Thermo Fisher Scientific, Waltham, MA) in a DMEM/F12 medium containing 10% of fetal bovine serum (FBS, Thermo Fisher Scientific, Waltham, MA) and 50 U/ml penicillin G (Sigma-Aldrich, St Louis, MO) unless indicated otherwise. Live imaging was started 4 hrs after incubating the explant attached fully to the coverslip in the tissue culture incubator. The imaging was conducted in a chamber maintained at 37°C and 5% CO2 on either an inverted PerkinElmer UltraVIEW Time Lapse Image Analysis System with an CMOS camera or a Nikon A1R HD confocal system on an Ni-E upright microscope with a GaAsP detector. For UltraVIEW, a 10x objective was used for the lower magnification images, and a 63x objective was used for time-lapse imaging (pixel size is 0.216 × 0.216 μm). For Nikon A1R, 25x objectives were used for both the low (pixel sizes are 0.48 × 0.48) and high magnification (pixel sizes 0.16 × 0.16 μm) time lapse recordings. In both settings, Z-stacks of 30-60 μm thickness with a 0.5 μm step were taken at each time frame with 10 minute-interval between frames. Live imaging was conducted up to 41 hours.

For the microtubule inhibition experiments, we first determined the dose of nocodazole (Sigma-Aldrich, St Louis, MO) to use. Two doses of nocodazole, 5 μM and 33 μM, which had been used in cochlear explants studies (Szarama et al., 2012, Shi et al., 2005) were tested on E13.5 utricular explants for 24 hrs. While explants treated with 5 μM nocodazole showed largely intact HCs with reduced tubulin signals from the cytoplasm, the utricle explants treated with 33 μM nocodazole showed reduced number of HCs, which looked unhealthy or apoptotic. Therefore, the dose of 5 μM was used for live imaging studies. Nocodazole in DMEM/F12 with 10% FBS or media for washing out was added to the utricular explants directly without disturbing the position of the explant under the microscope.

### AAV virus

The AAV2.7m8-CAG-Emx2-P2A-tdTomato (2.2×10^12^ genome copies/ml (GC/ml)) and AAV2.7m8-CAG-tdTomato (5.4×10^12^ GC/ml) were synthesized by Vector Biolabs, Inc. The expression of *Emx2* and *tdTomato* were driven by the CAG promoter. Four hrs after dissection, AAV was added to the E13.5 utricular explants to achieve the concentration of 4.0 × 10^10^ GC in 100 μl of culture medium containing DMEM/F12 with 2 % of FBS and 50 U/ml penicillin G. One hour later, 100 μl culture medium was added to the culture and after overnight incubation, the culture was washed several times before changing to DMEM/F12 containing 10 % FBS and 50 U/ml penicillin G.

### Whole mount immunostaining

Dissected utricles were attached to the Cell-Tak coated coverslip and then fixed with 4% paraformaldehyde in PBS at room temperature for 15 minutes. After fixation, the tissue attached to the coverslip was washed three times in PBS before blocking with PBS containing 5% donkey serum and 0.3% Triton-X for 45 minutes. For primary antibody, mouse anti-βII spectrin (1:500; BD Biosciences, San Jose, CA), rabbit anti-Arl13b (1:500; Proteintech, Rosemont, IL), mouse anti-acetylated tubulin (1:500; Sigma-Aldrich, St Louis, MO), rabbit anti-ninein, (1:500; Abcam, Cambridge, UK) or rabbit anti-γ-tubulin (1:500; Sigma-Aldrich, St Louis, MO) was used for overnight incubation at 4°C. Secondary antibodies of either Alexa Fluor 488/647 donkey anti-mouse IgG (Thermo Fisher Scientific, Waltham, MA) or Alexa Fluor donkey anti-rabbit 405/647 (Thermo Fisher Scientific, Waltham, MA) was used at 1:500 dilution for 15 minutes at room temperature. After extensive washing with PBS, samples were mounted in ProLong Gold Antifade (Thermo Fisher Scientific, Waltham, MA).

SiR-tubulin (Spirochrome, Switzerland) was added to the live utricular samples attached on the coverslip to achieve a final concentration of 1 μM in the culture medium containing 10% FBS in DMEM/F12 and incubated for 10 hours before imaging. For imaging of γ-tubulin, acetylated-tubulin, and ninein staining as well as SiR-tubulin, Airyscan imaging was conducted on a Zeiss LSM 780 with an Airyscan attachment (Carl Zeiss AG, Oberkochen, Germany) using a 63 × 1.4 NA oil objective lens. The acquired images were processed by Zeiss Zen Black software v2.1 for deconvolution.

### In situ hybridization

In situ hybridization was conducted as described previously (Morsli et al., 1998). Digoxin-labeled RNA probes were synthesized for Sox2 (Evsen et al., 2013) and Emx2 (Simeone et al., 1992) as described.

### Analysis

To correct for sample drifting during imaging over time, the 3-D time-lapse image was processed using ImageJ (Schneider et al., 2012) and the plugin of PoorMan3DReg (http://sybil.ece.ucsb.edu/pages/software.html) to re-align HCs to the stabilized x-y positions after semi-automatic adjustment of the z-positions by ImageJ macro (provided by Dr. Sho Ota), manual z-stack regulator. In some utricular images, GFP signals in tdTomato-labeled HCs were segmented by using the ImageJ macro (provided by Dr. Sho Ota), Centrin-detector, that is capable of isolating GFP signals overlapping with tdTomato expression in HCs depending on signal thresholding. The processed images were used for subsequent analyses. For tracking of centrioles and determination of HC apical center, we performed spot and cell tracking algorithm based on Imaris 9.5.0 (Bitplane, Zurich, Switzerland) and these tracking data were used to generate the 3D reconstructed videos.

The entry of centriole movements into Phase II was determined retroactively when the DC is consistently moving towards the peripheral direction where hair bundles should be established. The tdTomato expression level was calculated by the relative fluorescent intensities of the tdTomato in HCs subtracted from those of surrounding supporting cells without tdTomato expression. The distance between both centrioles were measured on the coordinate of the tracking data, and the average distances of both centrioles were calculated from all the time points during each of the phases in individual cells. For the migration speed, the difference of coordinates of each centriole compared to the center of the HC in one frame (10 min) was measured, and the average speed of each centriole was calculated from all the time points after HC center was determined. For statistics, students t-test was used with p-values of less than 0.5%, 0.1% and 0.01% indicated by *, ** and ***, respectively.

## Acknowledgements

We thank Dr. Sho Ota for writing the Fiji macro and providing helpful advice on live imaging data analyses, Kevin Isgrig and Dr. Wade Chen at NIDCD for advice on AAV2.7m8 virus and Dr. Elizabeth Driver at NIDCD for her help with live imaging. We are also grateful to Drs. Brian Galletta, Matt Hannaford and Nasser Rusan at National Heart Lung and Blood Institute and Drs. Elizabeth Driver, Inna Belyantseva and Ronald Petralia at NIDCD, and members of the Wu Lab for their critical review of the manuscript. We also want to thank Michael Mulheisen for technical support in conducting the in situ hybridization experiments.

---

**Figure Supplement 2a.**
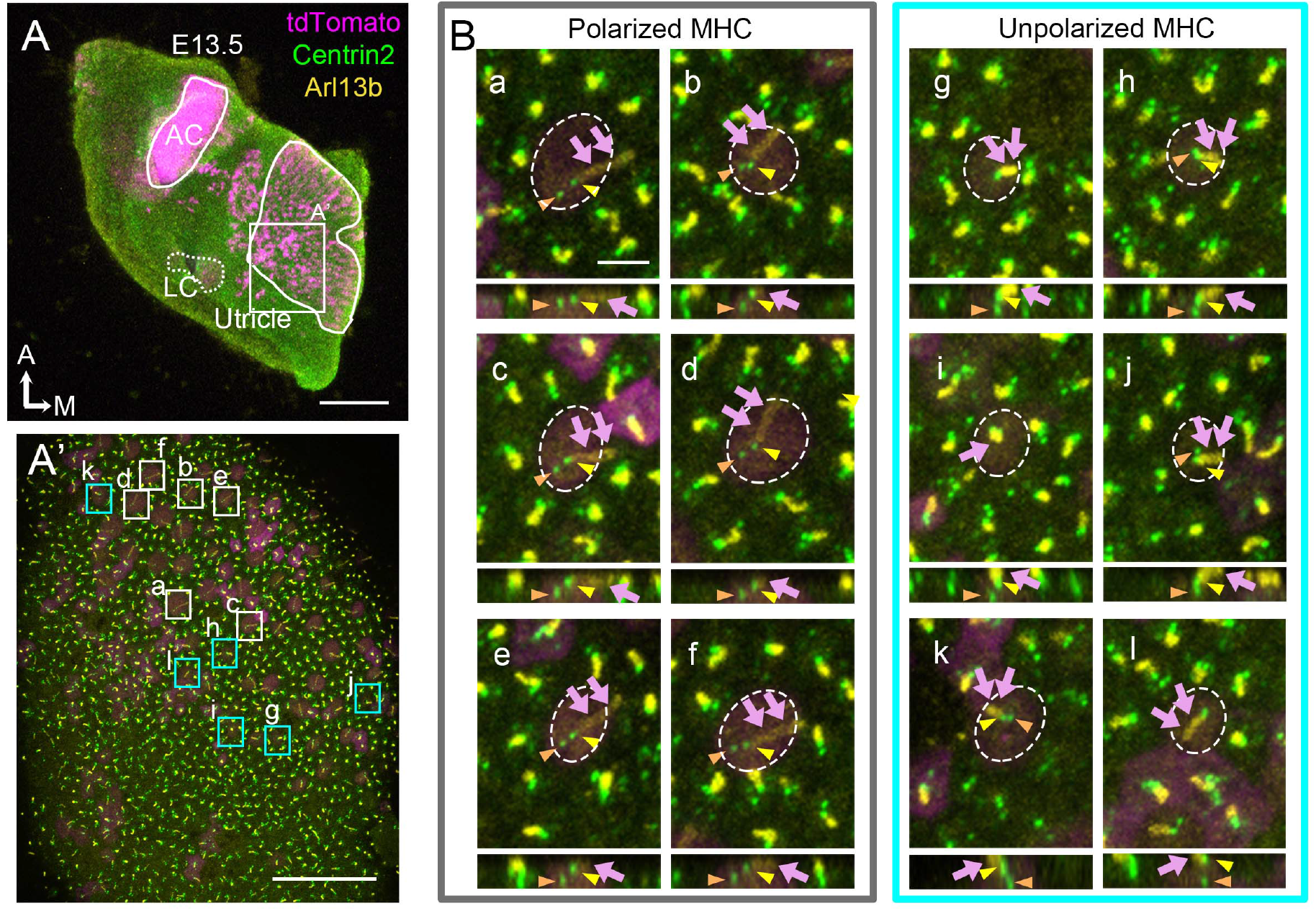
Identification of mother and daughter centrioles in polarized and unpolarized MHCs. (A,A’) A low (A) and high (A’) magnification of *Atoh1^Cre^*; *Rosa^tdT/+^*; *GFP-Centrin2* utricular explant that was immuno-stained with anti-Arl13b antibodies (yellow) at E13.5. (B) Representative apical and side views of polarized (a-f) and unpolarized (g-l) HCs in (A’) From apical views (upper panels), the MC (yellow arrowhead) is associated with the Arl13b-positive kinocilium (magenta arrows) and not the DC (orange arrowhead) in both polarized and unpolarized HCs. From side views (lower panels), the MC is positioned closer to the apical surface of the HC than the DC. Notably, the DC position is consistently more lateral than the MC in polarized HCs (a-f) but its position relative to the MC is variable in unpolarized HCs (g-l). Scale bars: 100 μm in (A), 30 μm in (A’) and 3 μm in (a), which applies to (b-l).

**Figure Supplement 2b.**
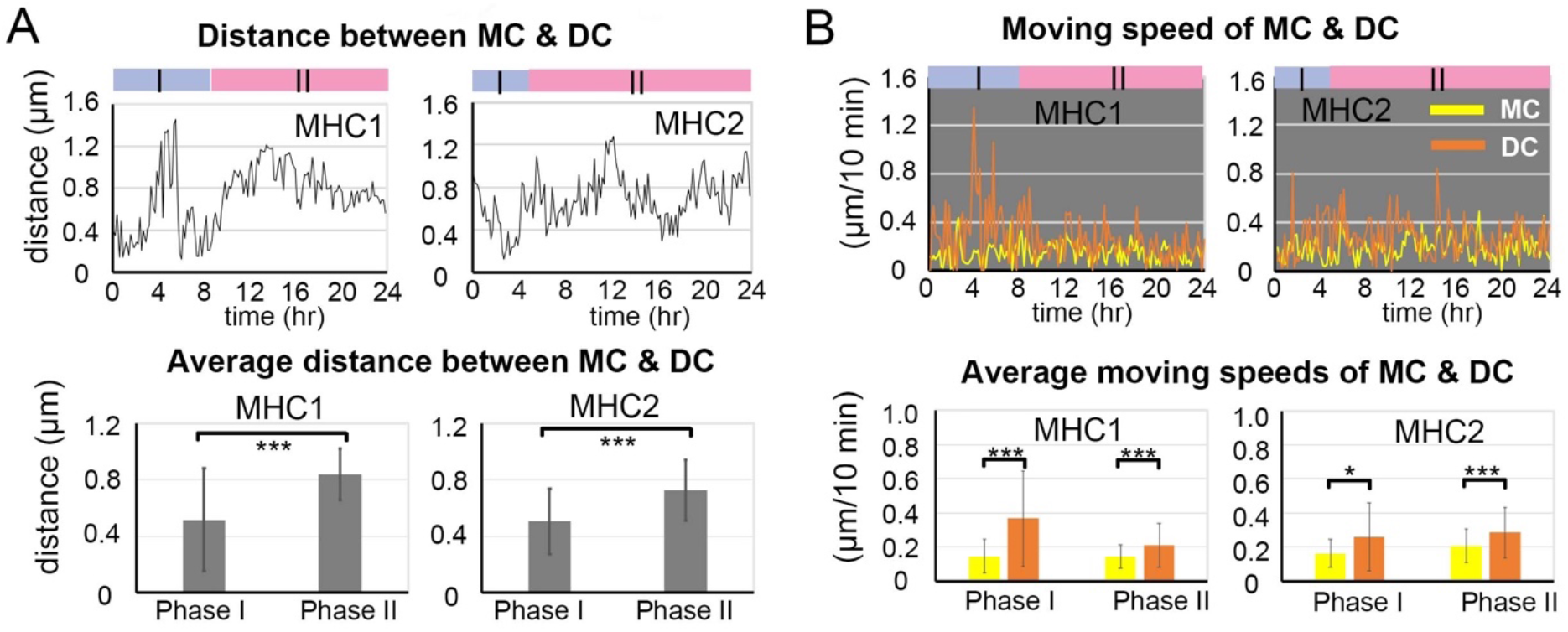
Traveled distance and speed between the MC and DC in MHCs. (A) The distance traveled between MC and DC over time and their average distances in the two phases. MHC1: number of time points measured equal (n) 53 and 92 for Phase I and II, MHC2: n=30 and 115 for Phase I and II. (B) Distance traveled by the MC and DC per a 10-min time frame and their average speed in the two phases. MHC1: number of speeds measured (n) equal 52 and 91 for Phase I and II, MHC2: n=29 and 114 for Phase I and II. Error bars represent standard deviation, SD. *p<0.05, *** P<0.001

**Figure Supplement 2c.**
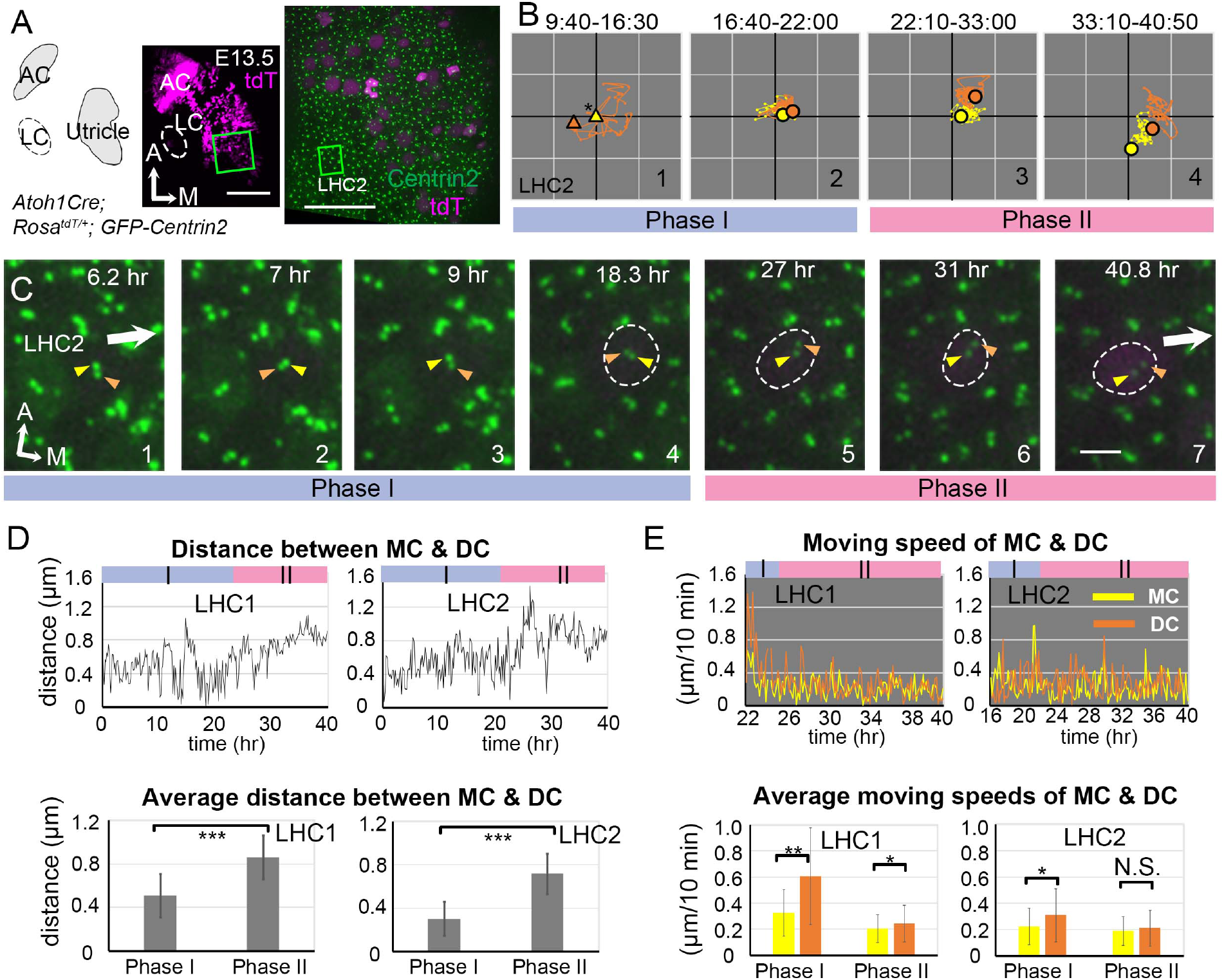
Live imaging of hair bundle establishment in LHCs based on centriole movements. (A) Schematic drawing and images of an *Atoh1^Cre^*; *Rosa^tdT/+^*; *GFP-Centrin2* utricular explant at E13.5. (B) The trajectory of the MC and DC in LHC2. The position of the MC (yellow triangle) is used as a proxy for the center of the HC (asterisk) for #1, because of the weak tdTomato signal. For #2-4, the center of the graph represents the apical center of the HC, as described in Fig. 2. The DC (orange) initially shows rapid movements around the MC in Phase I (#1-2). Then, the DC (orange) starts to move medially, which is followed by the MC in Phase II (#3-4). (C) Selected frames from a time-lapse recording of centriole movements showing apical views of LHC2. In Phase I (#1-4), the DC moves around the MC, when tdTomato signal is not apparent. At the end of Phase I (#4), the tdTomato signal is detectable and it continues to increase in Phase II. During Phase II (#5-7), the DC starts to migrate towards the medial periphery of the HC, which is followed by the MC. (D) The distance between MC and DC over time and their average distances in each phase. LHC1: number of time points measured (n) equal 146 and 100 for Phase I and II, LHC2: n=133 and 113 for Phase I and II. (E) The moving speed of MC and DC in each 10-min time frame and their averages in each phase. LHC1: number of speeds measured (n) equal 18 and 99 for Phase I and II, LHC2: n=34 and 112 for Phase I and II. Error bars represent SD in (D) and (E). * P<0.05, ** P<0.01, *** P<0.001.

**Figure Supplement 2d.**
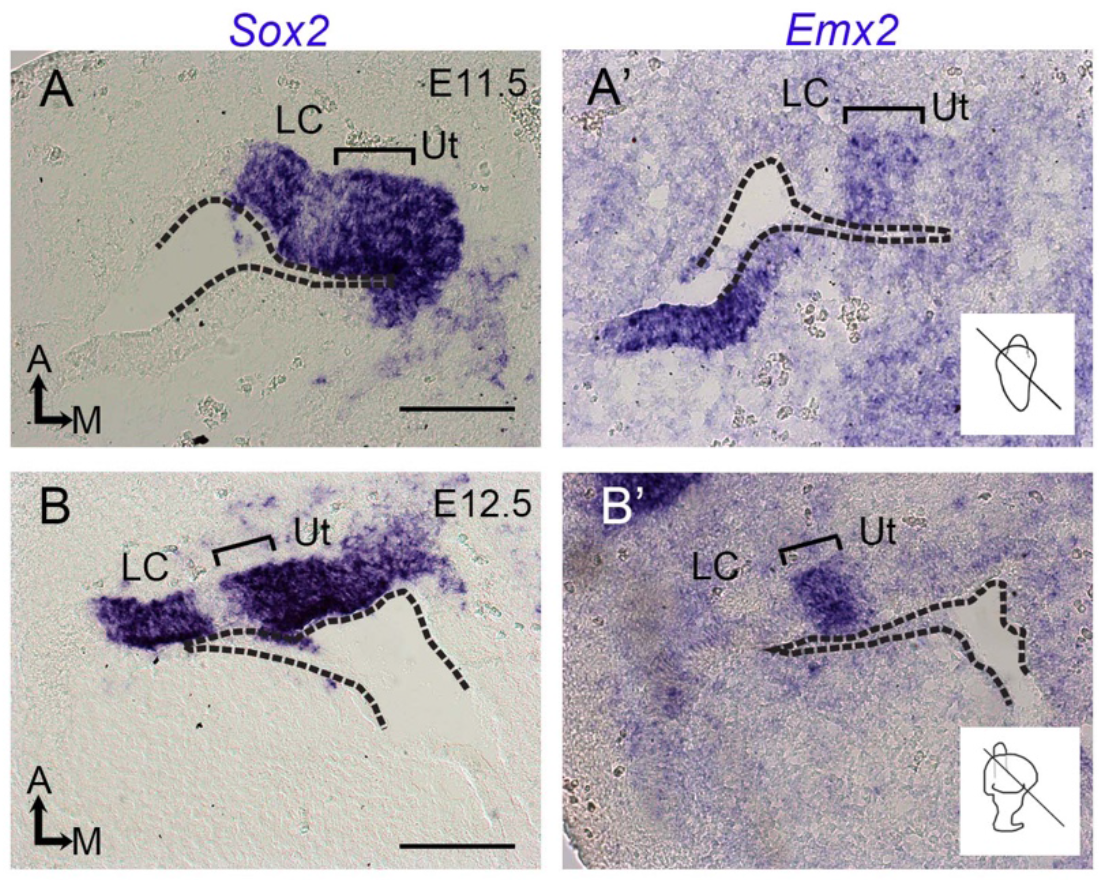
Expression of *Emx2* in the developing utricle. (A, A’, B, B’) In situ hybridization of adjacent sections at the levels of the utricle (Ut) and lateral crista (LC) at E11.5 (A, A’) and E12.5 (B, B’). The levels of sections are indicated in the inner ear schematics. Sensory tissues of the lateral crista and utricle are Sox2-positive, and the bracket indicates the lateral sensory region of the utricle that also expresses *Emx2*. Dotted lines indicate the apical margin of the otic epithelium. Scale bars: 50 μm.

**Figure Supplement 3a.**
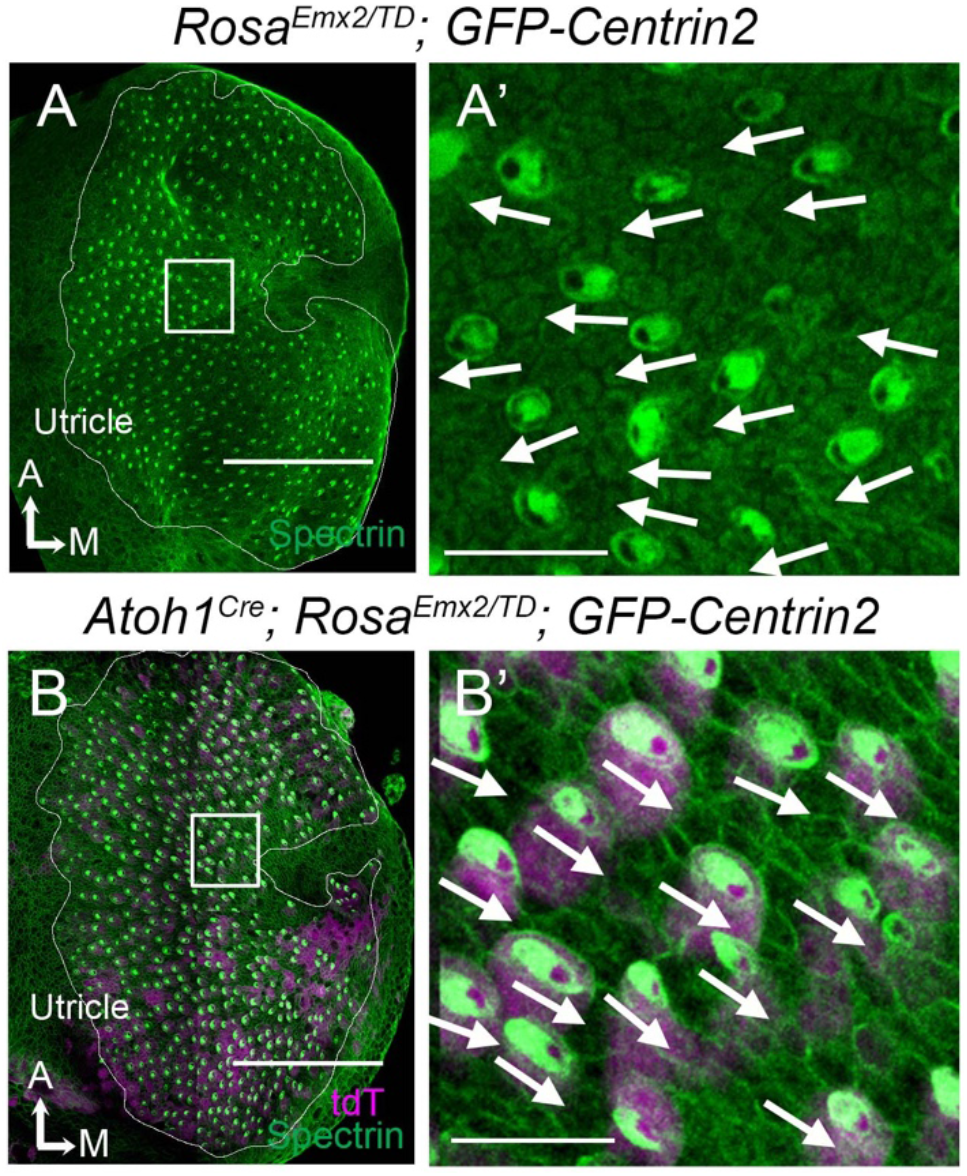
Gain-of-function of *Emx2* reverses hair bundle orientation in MHCs of *Atoh1^Cre^; Rosa^Emx2/tdT^; GFP-Centrin2* utricles. (A) Low magnification image of a *Rosa^Emx2/tdT^; GFP-Centrin2* utricle at E15.5. (A’) High magnification image of the rectangular area in (A), showing normal lateral-oriented hair bundles in the medial utricle (white arrows) based on β-spectrin immunostaining (green color). The void of β-spectrin staining indicates the position of the kinocilum and thus the orientation of the hair bundle. (B) Low magnification image of an *Atoh1^Cre^*; *Rosa^Emx2/tdT^*; *GFP-Centrin2* utricle at E15.5, in which HCs are tdTomato-positive (magenta) and stained with anti-β2 spectrin antibodies (green). (B’) High magnification of the rectangular area in (B) showing that hair bundles in the medial utricle all point towards the medial (white arrows) instead of the lateral direction as in controls (A’). The utricle is fan-shaped. Therefore, depending on selected regions, hair bundle orientation is not necessarily consistent among regions but always consistent within a region. Scale bars: 100 μm in (A, B) and 10 μm in (A’, B’).

**Figure Supplement 3b.**
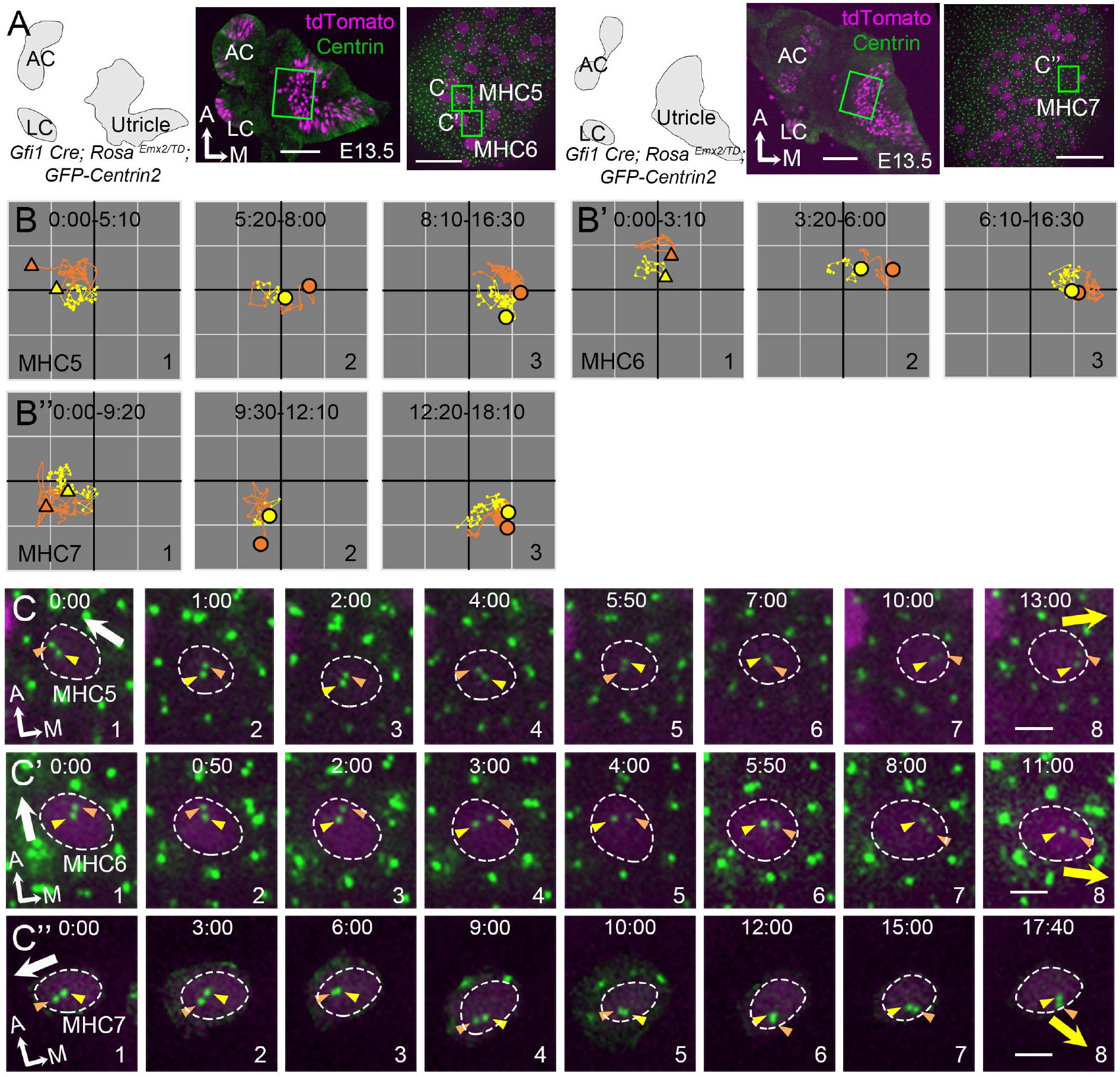
Centriole trajectories in *Emx2* Gain-of-Function MHCs using *Gfi1^Cre^*. (A) Schematic drawings, low and high magnification images of two *Gfi1^Cre^*; *Rosa^Emx2/tdT^*; *GFP-Centrin2* utricular explants. (B-B’’) The trajectory of MC and DC in MHC5 (B), MHC6 (B’), and MHC7 (B”). At first, both centrioles are positioned in the lateral side of the HC (B-B”, #1). Then, the daughter centriole starts to reverse its direction and moves towards the medial side of the HC (B-B”, #2), followed by the mother centriole (B-B”, #3). Yellow and orange triangles show the initial MC and DC positions in #1, whereas yellow and orange circled dots represent the final positions in Panels #2-#3. Each small grid is 1.25 × 1.25 μm. (C-C’’) Selected apical views of a live-imaging recording of MHC5 (C), MHC6 (C’) and MHC7 (C”) that overexpress *Emx2*. Initially, the DC (orange arrowhead) is located by the lateral side (white arrow, MHC5, MHC7 #1-4, MHC6 #1-3), then it changes course and moves to the medial side of the MC (yellow arrow, MHC5, 7 #7-8, MHC6, #4-8). Scale bars: 100 μm (low mag) and 30 μm (high mag) in (A), and 3 μm in (C), (C’) and (C’’).

**Video 1.**
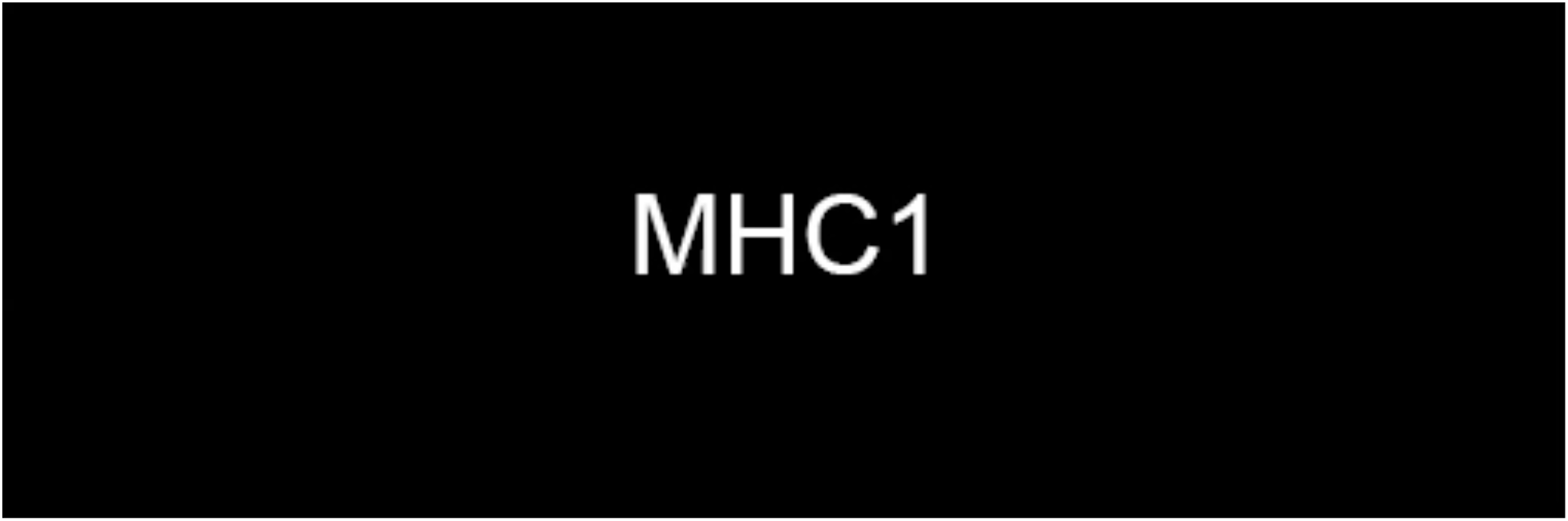
Time-lapse videos of the apical view (left panel) and the reconstructed image (right panel) of MHC1 and MHC2 in *Atoh1^Cre^; Rosa^tdT/+^; GFP-Centrin2* utricle showing the DC first moves sporadically around the MC. Then, the DC moves towards the lateral side (white arrow). This trajectory is followed by the MC. Yellow and red spheres indicate mother and daughter centrioles, respectively. Magenta indicates tdTomato signal within the HC. Scale bar: 3 μm. Play back speed: 7 frames per second.

**Video 2.**
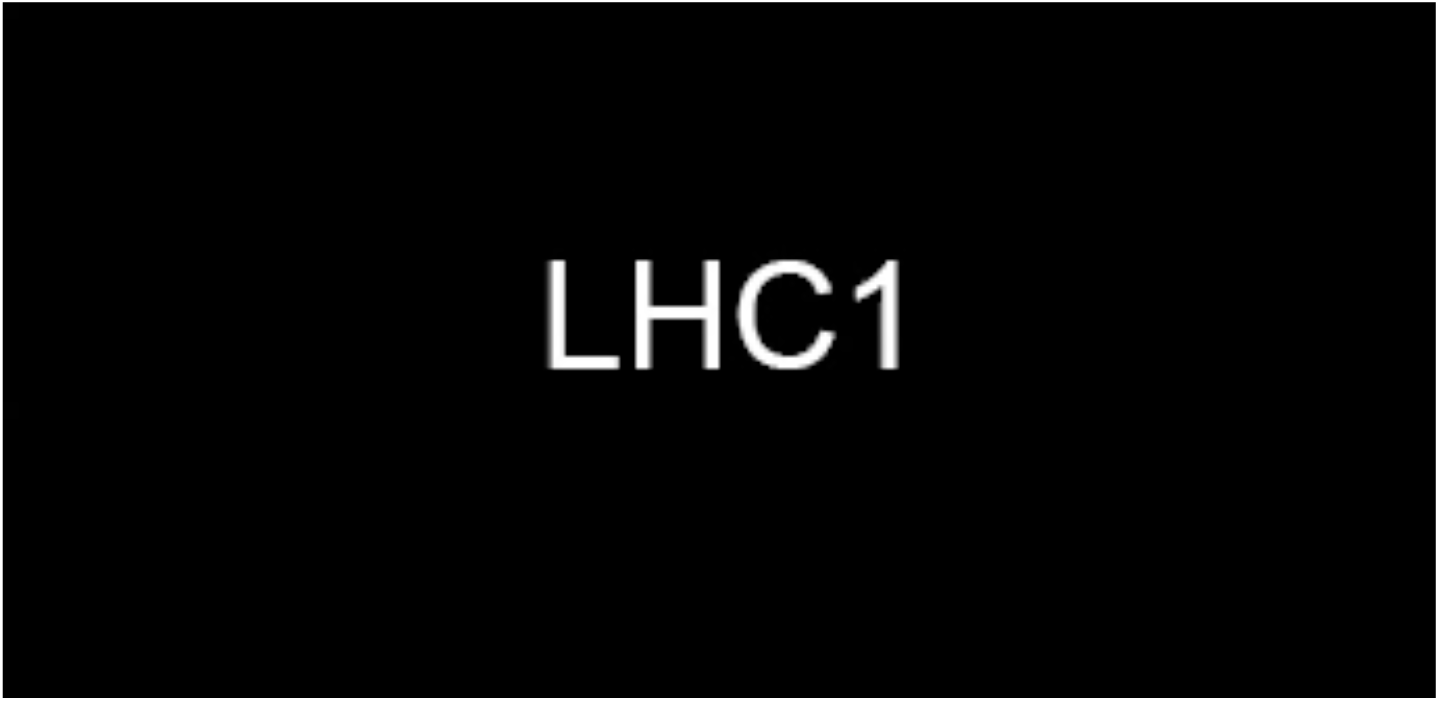
Time-lapse videos of the apical view (left panel) and the reconstructed image (right panel) of LHC1 and LHC2 in *Atoh1^Cre^; Rosa^tdT/+^; GFP-Centrin2* utricle, showing the DC first moves sporadically around the MC. Then, the DC moves towards the medial side (white arrow), which is followed by the MC. Yellow and red spheres indicate mother and daughter centrioles, respectively. Magenta indicates tdTomato signal within the HC. Scale bar: 3 μm. Play back speed: 7 frames per second.

**Video 3.**
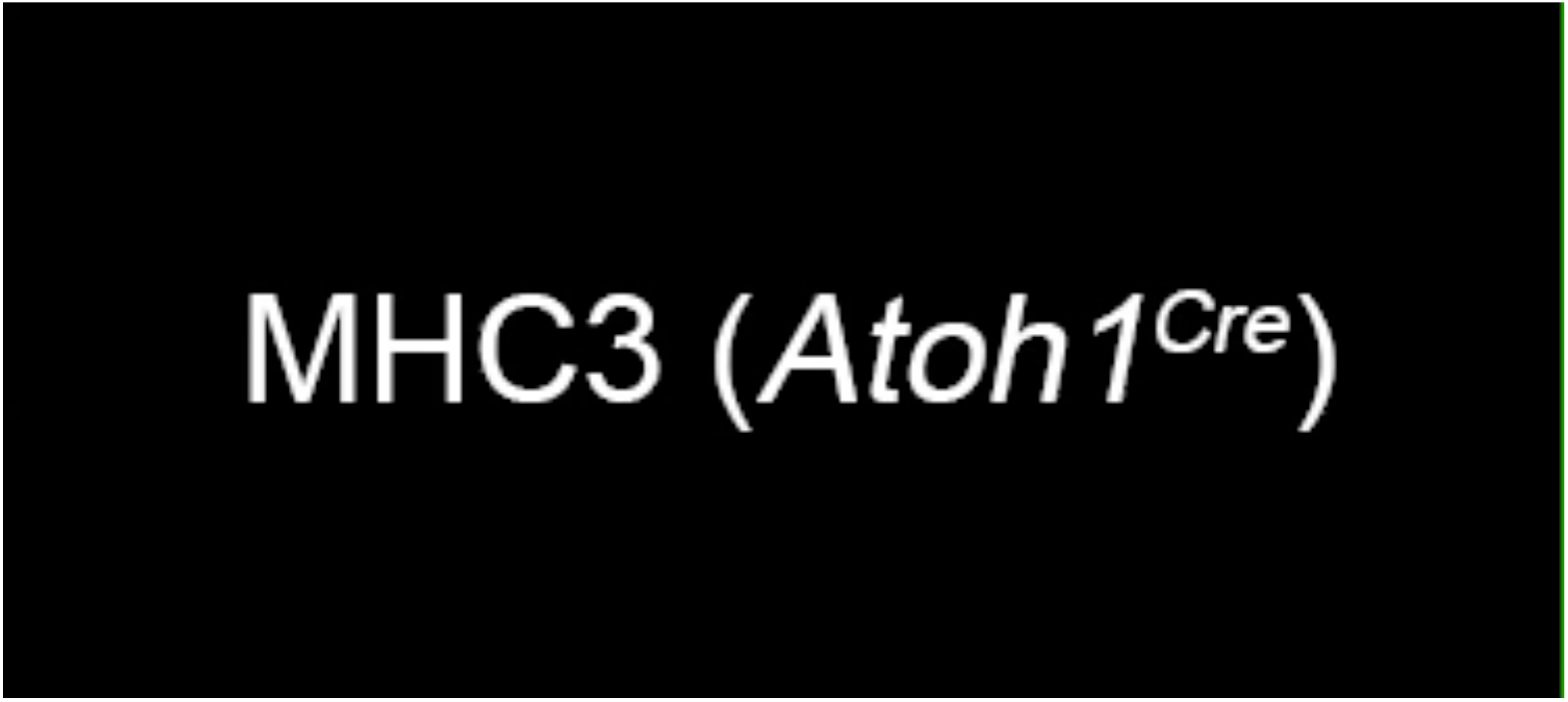
Time-lapse videos of the apical view (left panel) and the reconstructed image (right panel) of MHC3 (*Atoh1^Cre^; Rosa^Emx2/tdT^; GFP-Centrin2*) and MHC4 (*Gfi1^Cre^; Rosa^Emx2/tdT^; GFP-Centrin2*). The DC in MHC3 first moves sporadically around the MC, similar to other MHCs (video 1). Then, the DC moves towards the medial side (yellow arrow) in an opposite direction from normal MHCs (white arrow). This movement is followed by the MC. The DC in MHC4 first leads the MC moving towards the lateral side (white arrow) and then it changes course and moves medial to the MC towards the medial side (yellow arrow). Yellow and red spheres indicate mother and daughter centrioles, respectively. Magenta indicates tdTomato signal within the HC. Scale bar: 3 μm. Play back speed: 7 frames per second.

**Video 4.**
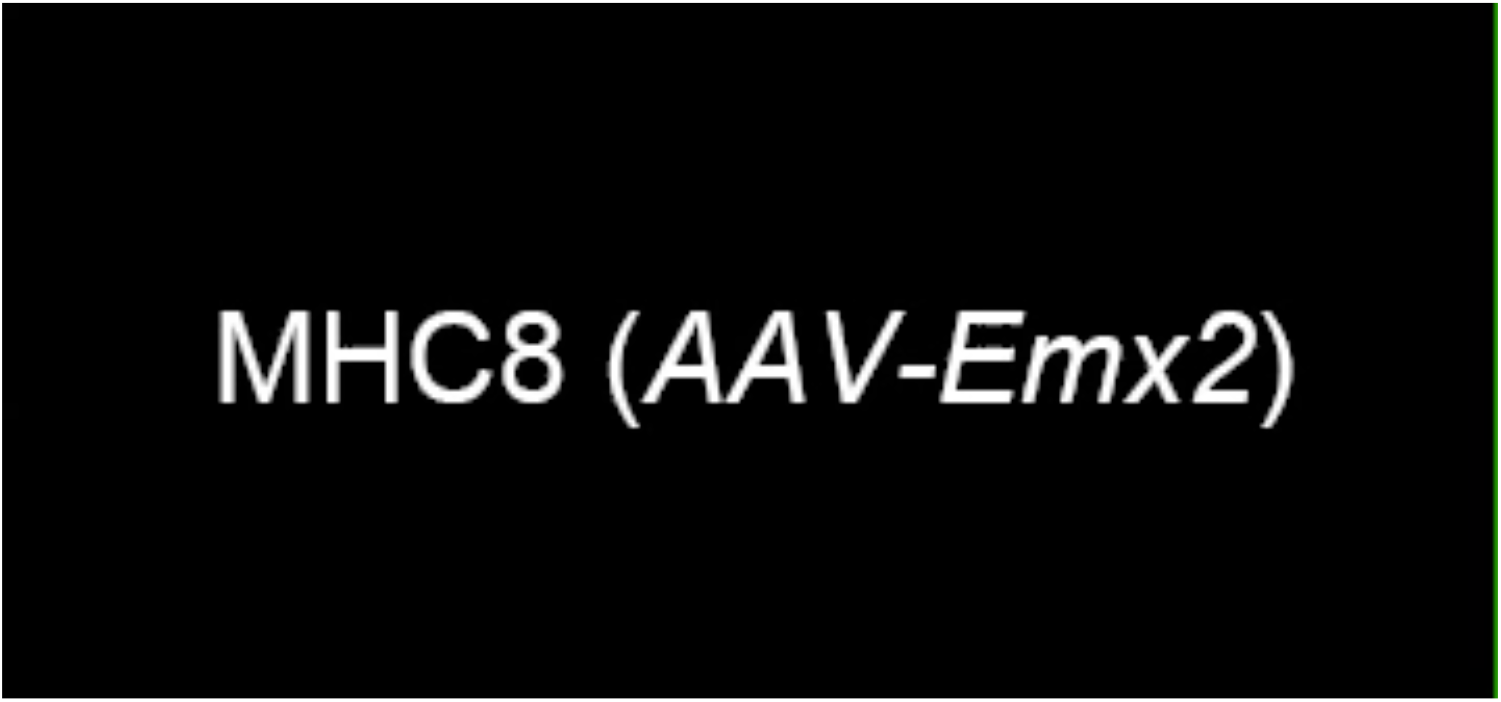
Time-lapse videos of the apical view (left panel) and the reconstructed image (right panel) of MHC8 and MHC9 in *GFP-Centrin2* utricular explant infected with AAV2.7m8-CAG-Emx2-P2A-tdTomato. In both MHC8 and MHC9, the DC starts out on the lateral side of the MC towards the lateral side of the utricle (white arrow). Then, it changes course to be on the medial side of the MC towards the medial utricle (yellow arrow). Yellow and red spheres indicate mother and daughter centrioles, respectively. Magenta indicates tdTomato signal in the infected HC cytoplasm. Scale bar: 3 μm. Play back speed: 7 frames per second.

**Video 5.**
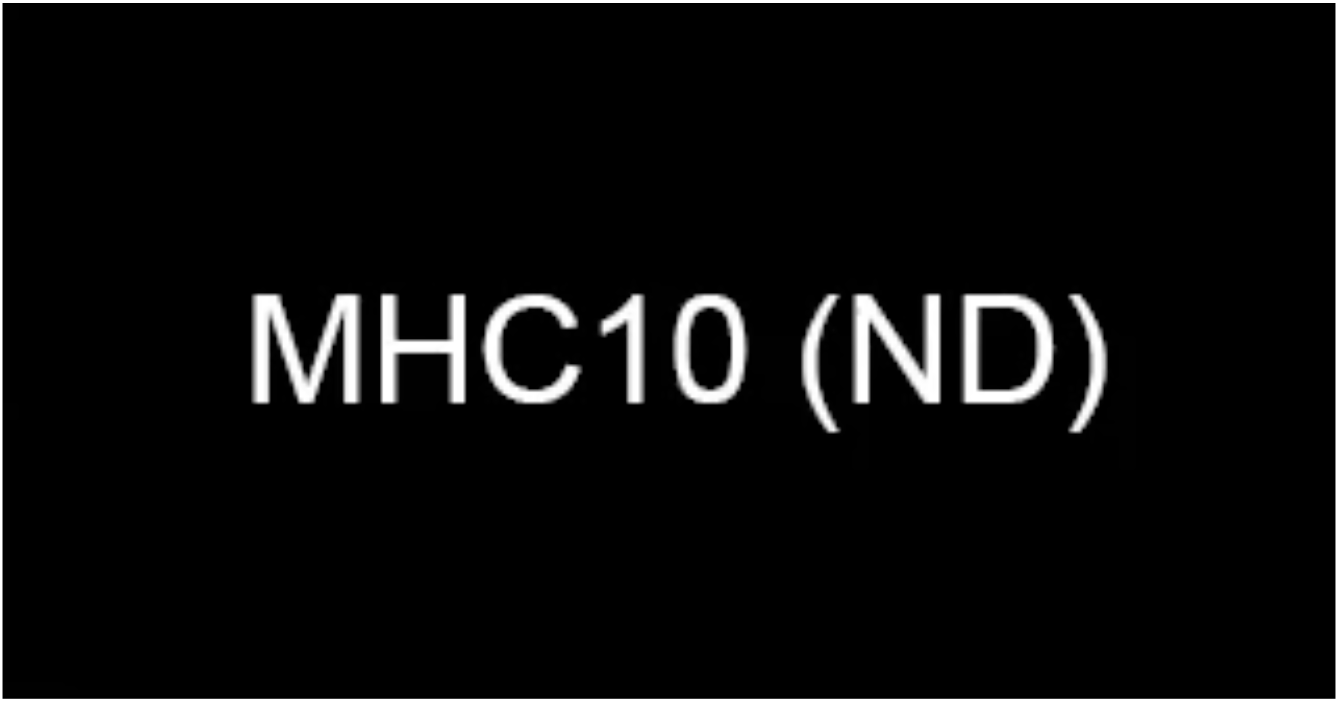
Time-lapse videos of the apical view (left panel) and the reconstructed image (right panel) of MHC 10 and MHC11 in *Atoh1^Cre^*; *Rosa^tdT/+^*; *GFP-Centrin2* utricle, showing that the introduction of nocodazole causes the lateral-positioned centrioles (white arrow) to spring back to the center of the HC and then they return to the peripheral position after nocodazole removal. Yellow and red spheres indicate mother and daughter centrioles, respectively. Magenta indicates tdTomato signal in the HC cytoplasm. Scale bar: 3 μm. Play back speed: 3 frames per second.

